# Longer metaphase and fewer chromosome segregation errors in modern human than Neandertal brain development

**DOI:** 10.1101/2022.03.30.486431

**Authors:** Felipe Mora-Bermúdez, Philipp Kanis, Dominik Macak, Jula Peters, Ronald Naumann, Mikhail Sarov, Sylke Winkler, Christina Eugster Oegema, Christiane Haffner, Lei Xing, Pauline Wimberger, Stephan Riesenberg, Tomislav Maricic, Wieland B. Huttner, Svante Pääbo

**Author notes:** These authors contributed equally; listed in alphabetical order (Philipp Kanis, Dominik Macak, Jula Peters). These authors jointly supervised the work; listed in alphabetical order (Wieland B. Huttner, Svante Pääbo).

## Abstract

Since the ancestors of modern humans separated from those of Neandertals, around one hundred amino acid substitutions spread to essentially all modern humans. The biological significance of these changes is largely unknown. Here, we examine all six such amino acid substitutions in the three proteins known to have key roles in kinetochore function and chromosome segregation and to be highly expressed in the stem cells of the developing neocortex. When we introduce these modern human-specific substitutions in the mouse, three substitutions in two of these proteins, KIF18a and KNL1, cause a prolongation of metaphase and a reduction in chromosome segregation errors in apical progenitors of the developing neocortex. Conversely, the ancestral substitutions cause a reduction in metaphase length and an increase in chromosome segregation errors in human brain organoids. Our data also show that, in these aspects, Neandertals were more similar to chimpanzees than to modern humans. Thus, the fidelity of chromosome segregation during neocortex development improved in modern humans after their divergence from Neandertals.

## Introduction

The neocortex is unique to mammals and the seat of sensory and motor activities(*1*). During the evolution of humans, the neocortex increased drastically in size. This is widely considered to be associated with the development of human cognitive abilities(*2*). Quantitative changes that lead to an increase in neocortex size include, for example, increases in the proliferative capacity and numbers of neocortical stem and progenitor cells, and consequently in the numbers of neurons and macroglial cells generated by them (*3–8*). Comparatively less is known about qualitative changes in neocortex development during hominin evolution that may have occurred concomitant with the increase in neocortex size. However, substitutions and duplications affecting the genes *FOXP2*(*9, 10*) and *SRGAP2C*(*11, 12*) have been shown to affect synapse formation and connectivity resulting in improved learning in mouse models(*13, 14*).

We previously compared the mitotic behavior of neocortical stem and progenitor cells in humans, chimpanzees and orang-utans, using induced pluripotent stem cell (iPSC)-derived cerebral organoids(*15*). We found that human proliferating apical progenitors (APs), the cells that line the ventricles and from which all other neural cells in the developing neocortex originate, spend around 50% more time in mitotic metaphase than the APs of chimpanzees and orang-utans. Metaphase is the step in mitosis where the cell finalizes the preparations to segregate and distribute the chromosomes equally to the two daughter cells(*16*). Hence, these differences in metaphase length raise the possibility that the fidelity of chromosome segregation during AP mitosis might differ between humans and apes, with potential consequences for neocortex development and function.

We focus on the roles of three proteins KIF18a (also known as (a.k.a.) Kinesin 8), KNL1 (a.k.a. CASC5), and SPAG5 (a.k.a. astrin), which are highly expressed in the germinal zones of the developing neocortex and are associated with mitotic spindle and kinetochore function. The kinetochore is a complex, three-dimensional, multi-protein structure mediating the attachment of chromosome centromeres with the ends of kinetochore microtubules(*17–19*).

An important role of kinetochores is to facilitate spindle assembly checkpoint (SAC) function that regulates the onset of chromosome segregation when chromosomes are correctly aligned at the metaphase plate(*20–23*). The three proteins stand out because they carry amino acid substitutions found in all present-day humans but are absent in apes as well as in Neandertals or Denisovans, *i.e*., so-called archaic humans, which separated from the evolutionary lineage leading to modern humans about half a million years ago(*24*). Any functional consequences of these substitutions would thus be unique to modern humans(*25, 26*).

KIF18a is a motor protein of the kinesin family that is involved in regulating correct chromosome positioning and attachment to kinetochore microtubules, and their bi-orientation within the mitotic spindle(*27–29*). KIF18a carries one modern human-specific amino acid substitution. KNL1 is part of the outer kinetochore, which is required for attachment of the kinetochores to the microtubules. It is also a main docking site for key proteins of the SAC, such as BubR1 and Mad1, and therefore important for chromosome alignment and segregation(*30–33*). KNL1 carries two modern human-specific amino acids. SPAG5 is a microtubule-associated protein recruited to kinetochores and is important for the stability of attachment of kinetochores to microtubules(*34–37*). SPAG5 carries three amino acid residues specific to modern humans.

We find that APs in the embryonic neocortex of mice where the modern human substitutions in KIF18a and KNL1 have been introduced by genome editing exhibit longer metaphases, more SAC-positive kinetochores, and fewer mis-segregating chromosomes. In converse experiments, where human embryonic stem cells that carry the ancestral variants of KIF18a and KNL1 are used to generate cerebral organoids, shorter metaphases, less SAC-positive kinetochores, and more chromosome segregation defects are observed in APs. Taken together, our data suggest that the three amino acid substitutions in KIF18a and KNL1 cause fewer chromosome inheritance errors to occur in APs of modern humans than in archaic humans and apes.

## Results

### More SAC-positive kinetochores in human than chimpanzee Aps

An active spindle assembly checkpoint (SAC), triggered, *e.g*., by one or more kinetochores not being properly attached to kinetochore microtubules, delay anaphase onset, and therefore prolong metaphase(*20, 38*). We first asked if induced pluripotent stem cell (iPSC)-derived cerebral organoids (below referred to as “organoids”) can serve as models to study SAC activity during neocortex development. As determined by immunofluorescence, the number of BubR1-positive kinetochores, a marker for active SACs, did not differ between APs that had formed a metaphase plate in human neocortex samples of gestation week (GW) 11-12, and those of human organoid APs of day 30-32 (13.6 vs. 12.5, p>0.05, Figure 1A, B, D). In agreement with this, another active SAC marker, Mad1, was also found to be present at kinetochores in similar numbers in the fetal and organoid APs (12.8 vs. 12.3, p>0.05, Supplementary Figure 1A, B, D). Organoids can therefore be used to study the number of SAC-positive kinetochores in cerebral tissue APs.

**Figure 1.**
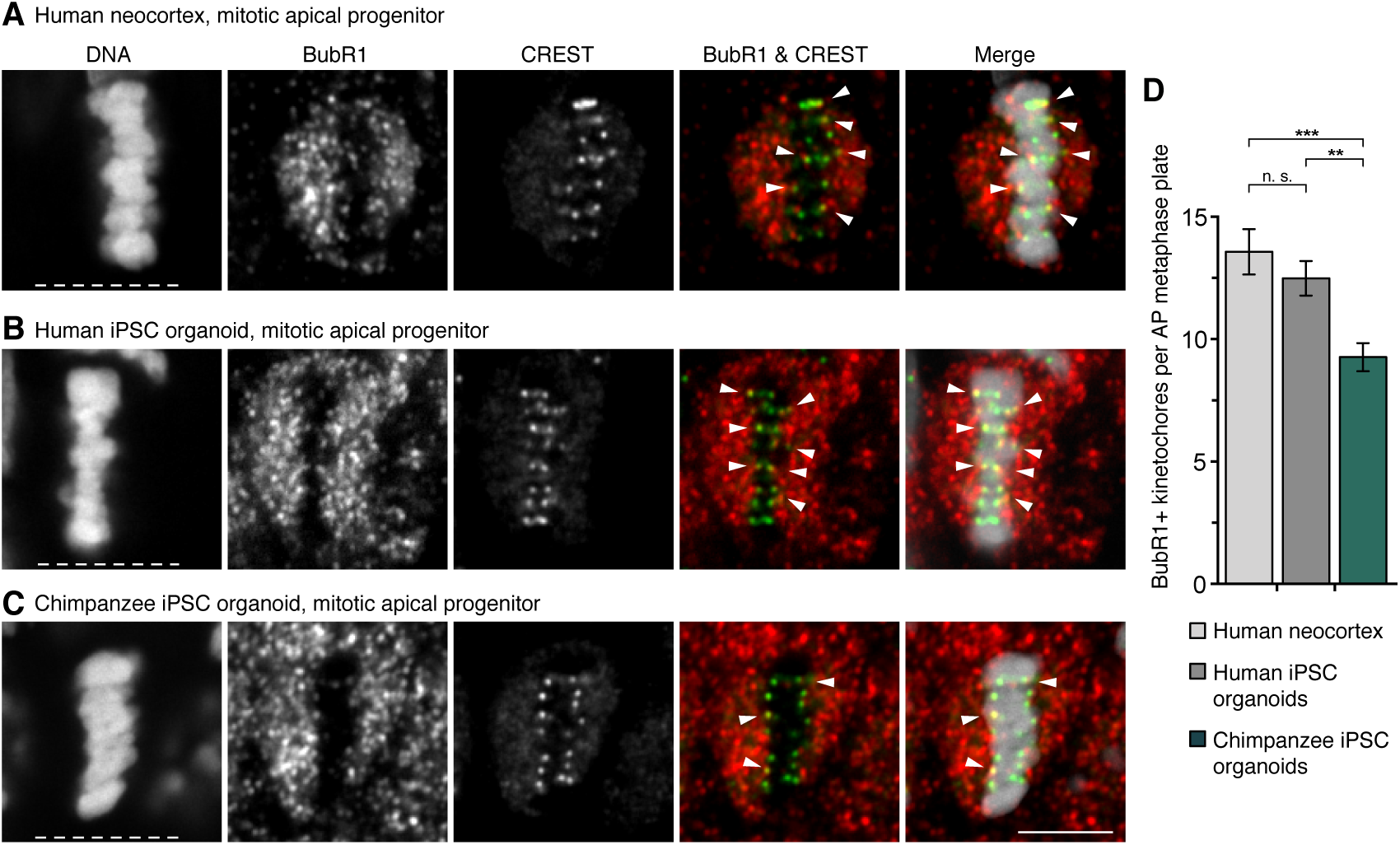
SAC-positive kinetochores in human and chimpanzee APs. (**A–C**) APs in metaphase stained with DAPI and immunostained for the SAC marker BubR1 (red in merges) and the kinetochore marker CREST (green in merges). Arrowheads indicate overlap of BubR1 and CREST immunoreactivity within the metaphase plate. (A) GW12 human neocortex; (B) Day 30 human iPSC-derived cerebral organoid; (C) Day 30 chimpanzee iPSC- derived cerebral organoid. White dashed lines, ventricular surface. Scale bar, 5 μm. (**D**) Quantification of kinetochores positive for BubR1 per AP metaphase plate, for the tissues in A–C (GW11-12, days 30-32). Data are the mean ± SEM of ≥41 APs from ≥3 independent experiments each, with 3 neocortex samples and ≥5 organoids each. Brackets with **, p <0.01; ***, p <0.001; n. s., non-significant (Kruskal-Wallis test with Dunn’s multiple comparisons correction).

To investigate if the previously described metaphase prolongation in human relative to chimpanzee APs(*15*) may be related to SAC activity, APs in human and chimpanzee iPSC-derived organoids were studied. The numbers of BubR1-positive kinetochores in fetal human neocortex and human organoid APs were 46% and 34% higher, respectively, than in chimpanzee organoid APs (p<0.001 and <0.01, respectively), Figure 1A-D). The numbers of Mad1-positive kinetochores were similarly 51% and 45% higher (both p<0.0001) in the fetal human neocortex and human organoid APs, respectively, than in chimpanzee organoid APs (Supplementary Figure 1A-D). Thus, human metaphase APs have more SAC-positive kinetochores than chimpanzee APs.

### Fewer lagging chromosomes in human than chimpanzee Aps

Differences in metaphase length and SAC signaling could affect chromosome segregation accuracy. To test this, the APs with chromosomes or large chromosome fragments lagging in anaphase and telophase in human and chimpanzee iPSC-derived organoids were quantified by serial confocal microscopy (Figure 2A–D). As expected for a process where mistakes need to be minimized to ensure normal physiological development, lagging chromosomes were rare (Figure 2E).

**Figure 2.**
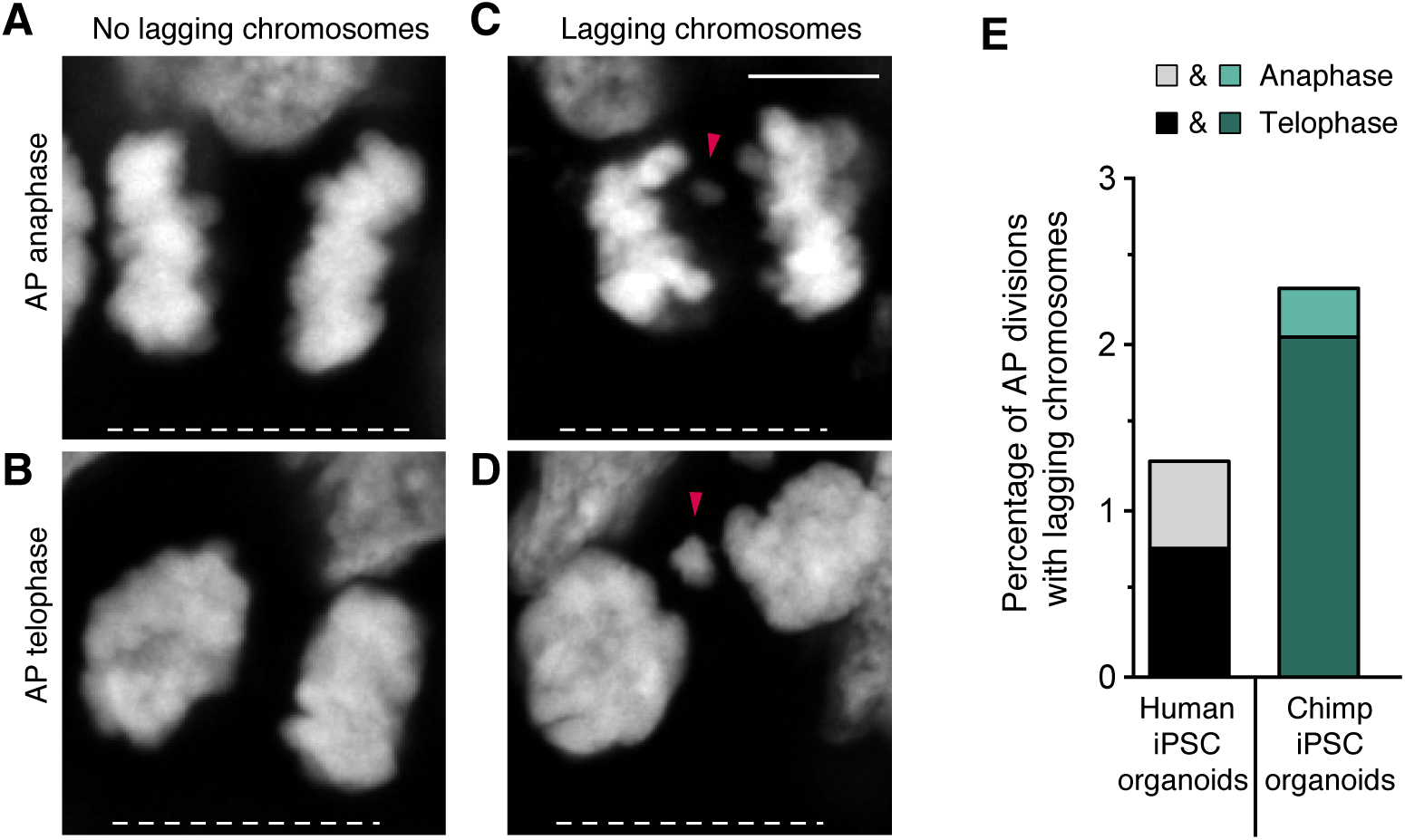
Lagging chromosomes in human and chimpanzee APs. (**A–D**) Mitotic apical progenitors (APs) in day 30-32 modern human iPSC-derived cerebral organoids stained with DAPI. Examples of APs in anaphase (A) and telophase (B) without lagging chromosomes, and in anaphase (C) and telophase (D) with lagging chromosomes, or large chromosome fragments (arrowheads). White dashed lines, ventricular surface. Scale bar, 5 μm. (**E**) Cumulative quantification (see Materials and Methods) of the percentages of AP divisions with lagging chromosomes in day 30-32 modern human (greyscale) and chimpanzee (Chimp, green) iPSC-derived cerebral organoids. Data are the sum from ≥5 independent experiments and a total of ≥342 AP divisions (only anaphase or telophase) each. Each bar shows the data for the sum of mitotic APs in anaphase plus telophase, with the stacked percentages for telophase (dark shade) and anaphase (light shade) being indicated separately.

Nevertheless, the percentage of mitotic APs with lagging chromosomes was higher in chimpanzee than human organoids (2.3% vs. 1.3%, or 8/342 in chimpanzee vs. 5/385 in human; Figure 2E). Interestingly, the majority of lagging chromosomes in both species was found in telophase rather than anaphase (10 out of 13 cells; Figure 2E), suggesting that lagging chromosomes are unlikely to be rescued by the end of cell division. Thus, AP chromosome segregation fidelity is higher in modern humans than in chimpanzees.

### Mice “humanized” for KIF18a, KNL1 and SPAG5

To ask if molecular changes unique to modern humans compared to not only apes but also Neandertals and Denisovans might lead to mitotic AP differences, we investigated six modern human-specific amino acid substations in KIF18a, KNL1 and SPAG5, three proteins involved in cell division and kinetochore function. In mice, where the amino acids at those six positions are identical to archaic humans and apes, we used CRISPR/Cas9 technology to change them to the modern human variants (“modern- humanized” mice, for simplicity referred to as “humanized” (h) mice, Figure 3A, B; see also Materials and Methods and Suppl. Figure 2 for generation and verification of the lines).

**Figure 3.**
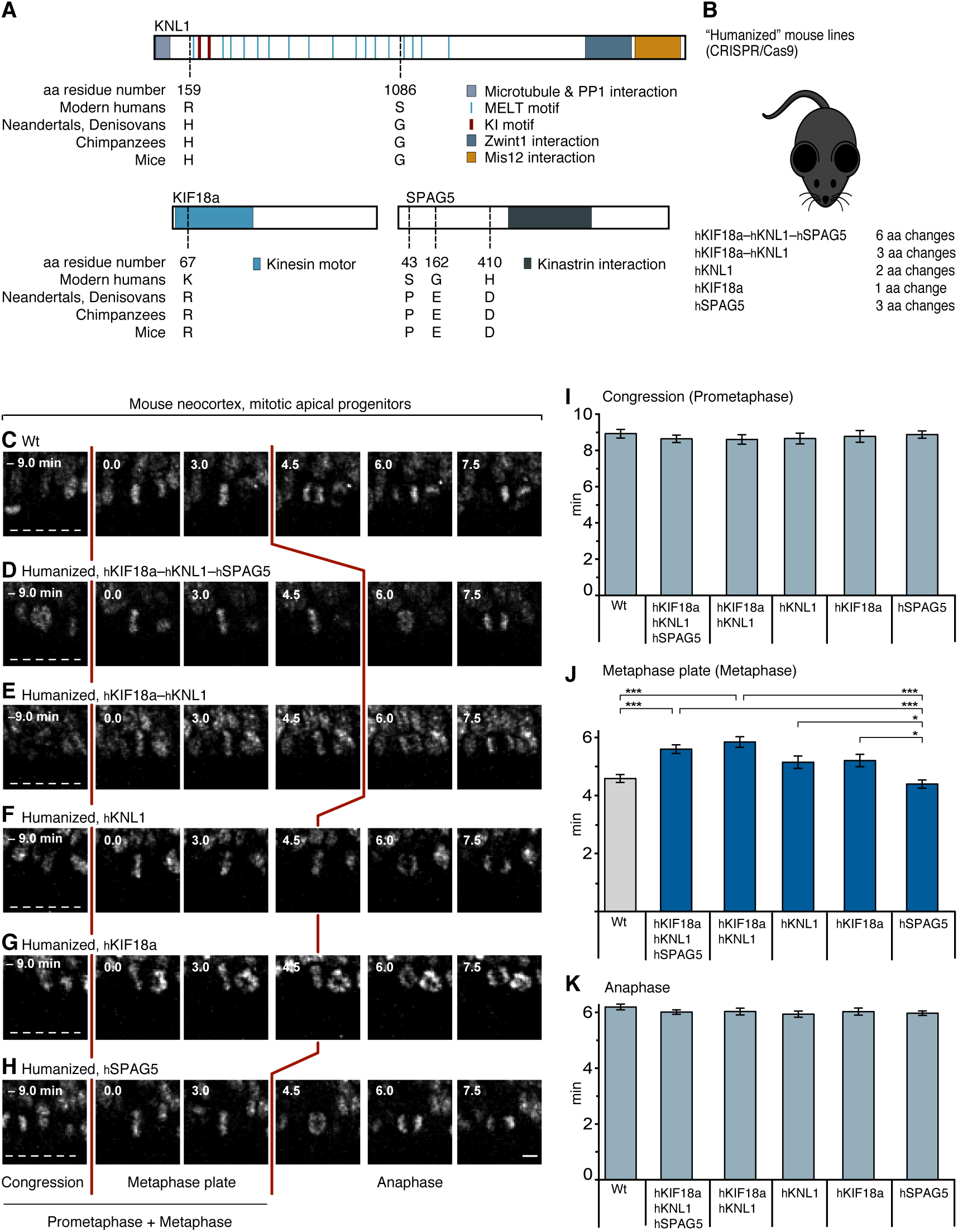
AP mitoses in mice humanized for Kif18a, Knl1 and Spag5. (**A**) Schematic representation of the domain structure of KNL1, KIF18a and SPAG5, carrying a total of six amino acid (aa) residue differences between modern humans (h) and archaic humans (a): KNL1, H_a_159R_h_ and G_a_1086S_h_; KIF18a, R_a_67K_h_; SPAG5, P_a_43S_h_, E_a_162G_h_ and D_a_410H_h_. Note that in wild type mice, these six aa residues are identical to those in archaic humans and apes. (**B**) C57Bl6/NCrl wt mice were gene-edited using CRISPR/Cas9 (see Materials and Methods) to change the wild type (Neandertal-like) aa residue(s) specified in (A) to the modern human variants. Five different humanized (h) mouse lines were generated as indicated. (**C-H**) Live-tissue imaging of the indicated mitotic phases of APs in organotypic coronal slice cultures of neocortex of E11.5 mice of the indicated genotypes (see Materials and Methods; wt, wild type). Zero (0) min is metaphase plate onset. Time-lapse intervals are 1.5 min. Red lines indicate the duration of metaphase, white dashed lines the ventricular surface. Scale bar, 5 μm. (**I**) Times between the start of chromosome congression and metaphase plate onset (referred to as “Congression” or “Prometaphase”); (**J**) between metaphase plate onset and chromatid segregation onset (referred to as “Metaphase plate” or “Metaphase”); and (**K**) between the onset of chromatid segregation and onset of chromosome decondensation (referred to as “Anaphase”), for APs in the six neocortical tissues described in (C–H). Data are the mean ± SEM of ≥50 APs from ≥3 independent experiments, with a total of ≥5 neocortices, for each of the six lines. Brackets with *, p <0.05; ***, p <0.001 (Kruskal- Wallis test with Dunn’s multiple comparisons correction).

First, a mouse line humanized for all six amino acids was examined, *i.e*. one in KIF18a (R67K), two in KNL1 (H159R and G1086S) and three in SPAG5 (P43S, E162G and D410H, Figure 3A), referred to as hKIF18a–hKNL1–hSPAG5 mice (Figure 3B). Live-tissue imaging of AP divisions in organotypic slice cultures of E11.5 hKIF18a–hKNL1–hSPAG5 neocortex showed an increase from 4.6 to 5.6 min in mean metaphase length relative to wild type mice (*i.e*. plus 22%, p<0.001, Figure 3C, D, J). To find out if substitutions in all three proteins participated in the metaphase prolongation, hKIF18a–hKNL1 mice humanized for the three amino acid substitutions in KNL1 and KIF18a were examined. These mice showed an increase in AP metaphase duration from 4.6 to 5.8 min (p<0.001,) that did not differ (p>0.05,) from the hKIF18a–hKNL1–hSPAG5 mice with the six amino acid substitutions (Figure 3C, E, J). This suggests that substitutions in both, or either of, hKNL1 and hKIF18a, but not hSPAG5, are involved in AP metaphase prolongation.

Next, three mouse lines were examined, each carrying one of the three genes in a humanized form. APs in the hKNL1 as well as hKIF18a lines showed a tendency toward longer metaphases compared to the wt (by 11% and 13%, respectively, or 5.1 and 5.2 min instead of 4.6 min) albeit not significantly so (p>0.05,) (Figure 3F, G, J). This suggests that substitutions in hKNL1 and hKIF18a additively contribute to the AP metaphase prolongation. In contrast, in hSPAG5 mice, the length of the AP metaphase was 4.4 min and hence similar to wild type mice (Figure 3H, J), indicating that the substitutions in hSPAG5 are not involved in the metaphase prolongation in the hKIF18a–hKNL1–hSPAG5 mice. It also shows that humanization of any three amino acids in a protein involved in mitosis and kinetochore function is not sufficient to prolong AP metaphase.

Notably, the durations of prometaphase, which precedes metaphase, and of anaphase, which follows metaphase, did not show any significant changes in any of the mouse lines analyzed (Figure 3I, K). Together, these results suggest a specific role of the amino acid substitutions in KNL1 and KIF18a in the prolongation of AP metaphase seen in modern humans.

### Human organoids “ancestralized” for *KIF18a* and *KNL*1

To investigate if the amino acid substitutions in KIF18a and KNL1 prolong metaphase in a modern human genetic background, we used CRISPR technology to change the relevant single codon in *KIF18a* and the two codons in *KNL1* back to the ancestral, Neandertal-like state in the H9 embryonic stem cell (ESC) line. Individual cells subjected to the relevant RNA guides and donor DNAs were expanded to cell lines, among which we selected two independent “ancestralized” lines (aKif18a–aKNL1 lines 1 (L1) and 2 (L2)) and two lines where none of the three codons had been changed (“control” lines 1 (L1) and 2 (L2), Figure 4A) based on sequencing across the edited sites. To exclude mono-allelic deletions(*39*) and loss of heterozygosity at the CRISPR target sites as well as big chromosomal deletions or duplications elsewhere in the genome, the four lines were characterized by quantitative PCR of the target sites, by genotyping of heterozygous single nucleotide polymorphisms up- and down-stream of the sites, as well as by low coverage whole-genome sequencing. Furthermore, mRNA expression of *KIF18a* and *KNL1* was found to be similar among all four lines and similar to the expression in the cell line from which the cells were derived from (Supplementary Table 2).

**Figure 4.**
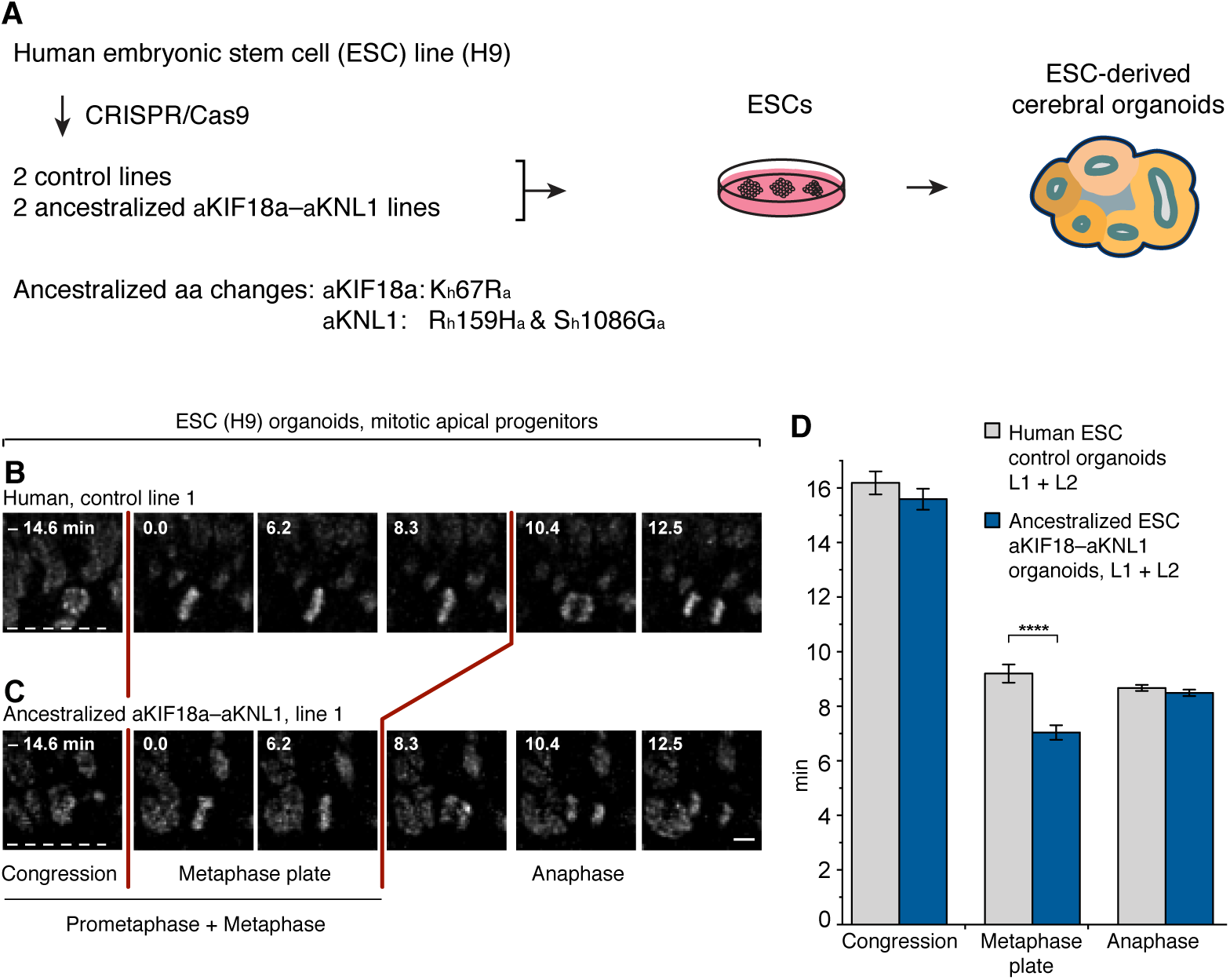
AP mitoses in organoids ancestralized for KIF18a and KNL1. (**A**) Human H9 embryonic stem cells (ESCs) were gene-edited using CRISPR technology to convert the indicated three amino acids in KIF18a and KNL1 from the modern human (h) to the ancestral (a) Neandertal-like states, and used to grow cerebral organoids (see Materials and Methods). (**B, C**) Live-tissue imaging of the indicated mitotic phases of APs in organotypic slice cultures of day 27-30 cerebral organoids grown from the indicated control (B) and ancestralized (C) line. Zero (0) min is metaphase plate onset. Time-lapse intervals are 2.08 min. Red lines indicate the duration of metaphase, white dashed lines the ventricular surface. Scale bar, 5 μm. (**D**) Times between the start of chromosome congression and metaphase plate onset (referred to as “Congression”), between metaphase plate onset and chromatid segregation onset (referred to as “Metaphase plate”), and between chromatid segregation onset and general chromosome decondensation onset (referred to as “Anaphase”), for APs in the two types of organoids as described for lines 1 in (B, C). Data are the mean ± SEM for organoids grown from the two control and the two ancestralized lines (L1, L2), and comprise ≥119 APs from four independent experiments, with ≥7 organoids, for each of the lines. Bracket with ****, p <0.0001 (Mann-Whitney U test).

Organoids were generated from the four lines, and live-tissue imaging of AP divisions in organotypic slice cultures was performed after 27-30 days of culture. The average metaphase lengths of APs in the organoids grown from the two ancestralized lines was 7.0 min, whereas it was 9.2 min for the control lines, *i.e*. a 24% decrease upon ancestralization (p<0.0001).

These results did not depend on any single control or aKIF18a–aKNL1 line, as AP metaphases were shorter in each of the aKIF18a–aKNL1 lines than in the control lines and the original non-edited ESC line (Supplementary Figure 3). Similar to the situation in the humanized mice, the durations of AP prometaphase and anaphase were not affected in any of the organoids (Figure 4B–D, Supplementary Figure 3E, F).

In conclusion, the mice and the human organoids that are “humanized” and “ancestralized”, respectively, for KIF18a and KNL1 show that the three modern human-specific amino acid substitutions in these two proteins specifically prolong metaphase in APs.

### Metaphase prolongation arises during cerebral cortical development

Our previous work(*15*) demonstrated that the increased metaphase duration in human APs relative to chimpanzee and orang-utan APs occur in organoids grown for approximately 30 days (as in the present study) but not in non-neural cells, including the iPSCs used to generate the organoids, nor in organoids at 52 days. It thus seems specific to early stages of cerebral cortical development in humans. To test if this holds true for the metaphase prolongation caused by the modern human-specific substitutions, the durations of mitotic phases were measured in monolayer cultures of the H9 ESCs used to generate the organoids. No differences in the duration of metaphase (Supplementary Figure 4A–E) or other mitotic phases (Supplementary Figure 4A–D, F, G) were observed between aKIF18a–aKNL1 and control H9 ESCs. Thus, the prolongation of AP metaphase in ancestralized KIF18a–KNL1 human organoids arises during the early development of the organoid tissue, similar to what is observed in human and chimpanzee organoids.

### SAC-positive kinetochores in humanized mice and ancestralized organoids

To test if the prolongation of AP metaphase in the mice humanized for KIF18a and KNL1 is associated with a change in the number of SAC-positive kinetochores, E11.5 mouse neocortices were immunostained for BubR1. In metaphase APs of mice humanized for KIF18a and KNL1, the average number of BubR1-positive kinetochores was 7.1, whereas it was 5.5 in wild type mice, *i.e*. a 29% increase upon humanization (p<0.01) (Figure 5A–C). In agreement with this, the number of Mad1-positive kinetochores in the humanized mouse APs was 7.5, whereas it was 5.5 in wild type mice, *i.e*. a 36% increase (p<0.01) (Supplementary Figure 5A–C).

**Figure 5.**
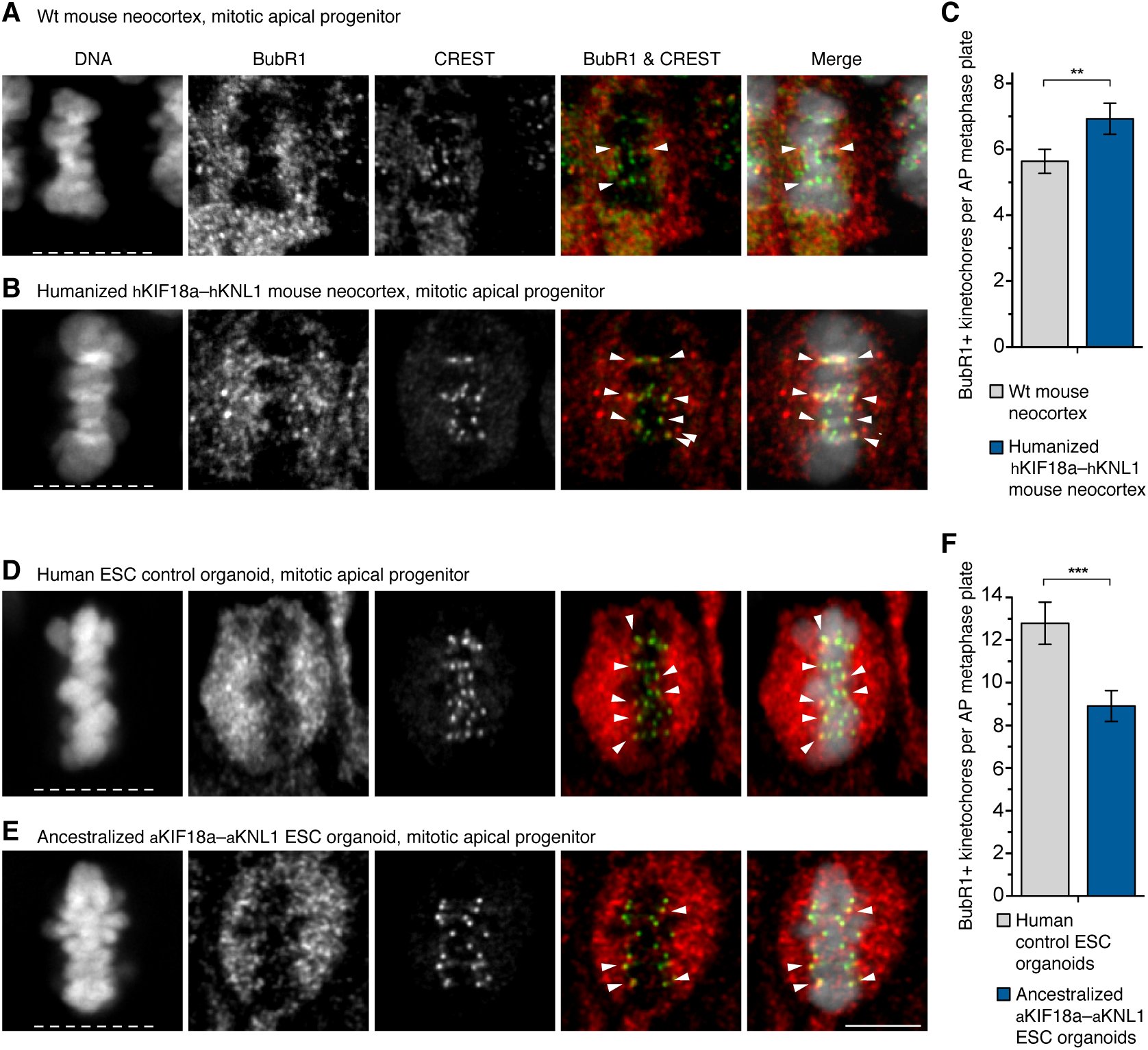
SAC-positive kinetochores in mice humanized for Kif18a and Knl1 and organoids ancestralized for KIF18a and KNL1. (**A, B, D, E**) APs in metaphase stained with DAPI and immunostained for BubR1 (red in merges) and the kinetochore marker CREST (green in merges). Arrowheads indicate overlap of BubR1 and CREST immunoreactivity within the metaphase plate. (**A**) Neocortex of E11.5 wild type (wt, ancestral-like) mouse; (**B**) neocortex of E11.5 mouse humanized for *Kif18a* and *Knl1*; (**D**) modern human non-edited day 27 cerebral organoid (control line 1); (**E**) day 29 organoid ancestralized for *KIF18a* and *KNL1* (edited line 1). White dashed lines, ventricular surface. Scale bar, 5 μm. (**C**) Number of BubR1-positive kinetochores per AP metaphase plate in mouse neocortex as described in (A, B). Data are the mean ± SEM of ≥41 APs from three independent experiments, with a total of ≥5 neocortices, for each of the two types of mice. Bracket with **, p <0.01 (Mann-Whitney U test). (**F**) Number of BubR1-positive kinetochores per AP metaphase plate for day 27-30 organoids (e.g., D, E). Data are the mean ± SEM of ≥54 APs from six independent experiments, with a total of ≥10 organoids, for each of the two ESC lines. Bracket with ***, p <0.001 (Mann-Whitney U test).

Similarly, APs in human ESC-derived organoids ancestralized for KIF18a and KNL1 were examined. They had an average of 8.9 BubR1-positive kinetochores whereas the control organoids had an average of 12.8 BubR1-positive kinetochores, *i.e*. a 30% decrease upon ancestralization (p<0.001; Figure 5D–F). Corresponding numbers for MAD1-positive kinetochores were 7.5 and 10.4, respectively, *i.e*. a 28% decrease (p<0.0001; Supplementary Figure 5D–F). Of note, the APs analyzed were all positive for FoxG1, indicating cortical or telencephalic identity (Supplementary Figure 5G). Thus, the number of SAC-positive kinetochores in APs increase in mice humanized for KIF18a and KNL1 while it decreases in human organoids ancestralized for these proteins.

### Lagging chromosomes in humanized mice and ancestralized organoids

To test if the persisting SAC activity and the delayed anaphase onset might lower the incidence of chromosomal segregation errors, the APs showing lagging chromosomes were quantified in anaphase or telophase in the neocortex of E11.5 mice humanized for KIF18a and KNL1 (Figure 6A-D). As for modern human and chimpanzee organoids (Figure 2E), few mitotic APs had lagging chromosomes (Figure 6I). However, the proportion of lagging chromosomes in wild type mice was almost twice that of the mice humanized for KIF18a and KNL1 (7/323 vs. 4/358) (Figure 6I). Similar to the situation in the modern human and chimpanzee organoids (Figure 2E), the majority of the mitotic APs with lagging chromosomes (8 of 11) were seen in telophase rather than anaphase, suggesting that most of these segregation defects do not become rescued by the end of cell division.

**Figure 6.**
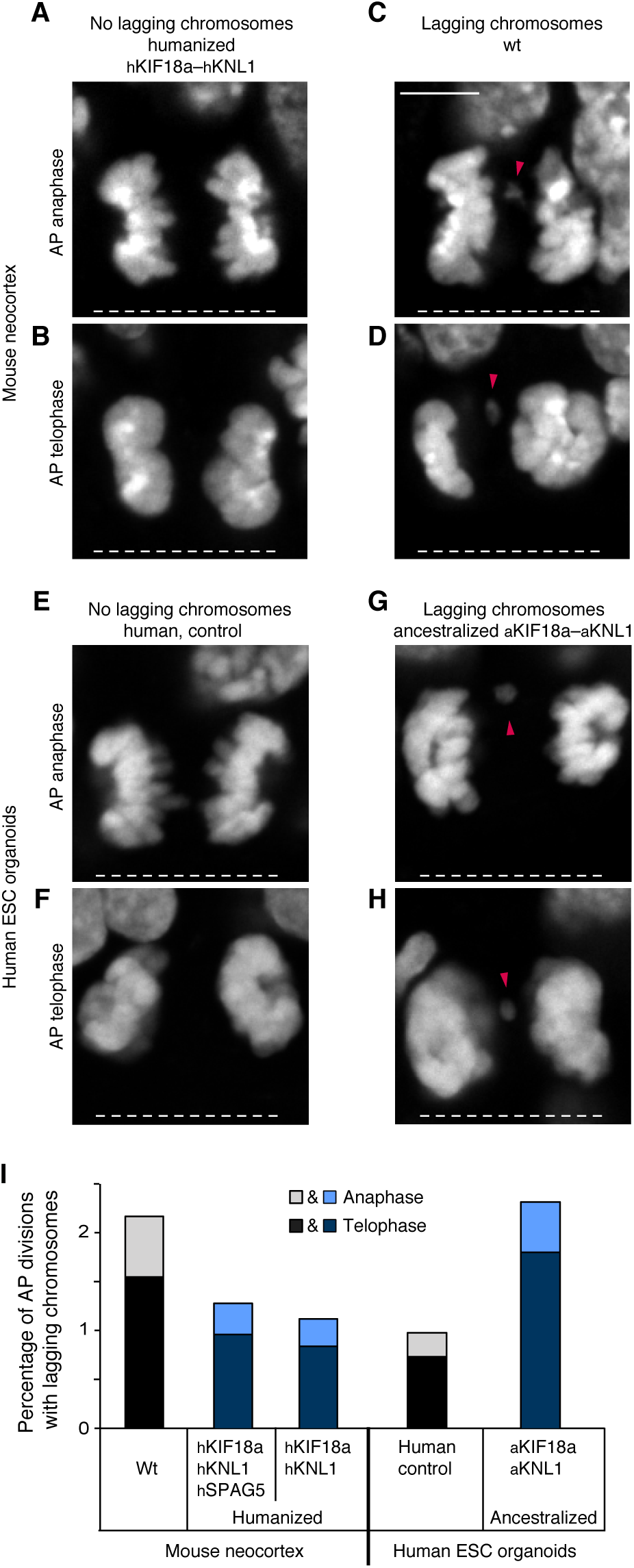
Lagging chromosomes in mice humanized for Kif18a and Knl1 and organoids ancestralized for KIF18a and KNL1. (**A–H**) Mitotic APs stained with DAPI in the neocortex of E11.5 wild type (wt) mice and mice carrying humanized *hKif18a–hKnl1* (A-D) and in day 27-30 control cerebral organoids (line 2) and organoids with ancestralized *aKIF18a–aKNL1* (line 2, E-H). Examples of APs in anaphase (A, E) and telophase (B, F) without lagging chromosomes, and in anaphase (C, G) and telophase (D, H) with lagging chromosomes, or large chromosome fragments (arrowheads). White dashed lines, ventricular surface. Scale bar, 5 μm. (**I**) Cumulative quantification (see Materials and Methods) of the percentages of AP divisions with lagging chromosomes for the tissues described in A–H, as well as for the neocortex of E11.5 mice carrying humanized *hKif18a–hKnl1–Spag5*. The mouse data are the sum of ≥314 AP divisions in anaphase or telophase from six experiments for each of the three types of mouse neocortex, the human organoid data are the sum of ≥389 AP divisions from six experiments for each of the two types of organoids (sums of lines 1 and 2). The percentages for telophase (dark shade) and anaphase (light shade) are indicated separately.

Given the small numbers of observations of lagging chromosomes, lagging chromosomes were also quantified in the mice humanized for SPAG5 as well as KIF18a and KNL1. The proportion of lagging chromosomes in these mice was 4/314, similar to that observed in the mice humanized for KIF18a and KNL1 (Figure 6I).

We similarly analyzed the fidelity of chromosomal segregation in the organoids ancestralized for KIF18a and KNL1. The proportion of mitotic APs with lagging chromosomes in the organoids ancestralized for KIF18a and KNL1 was more than twice than that in the control organoids (9/389 vs. 4/410, Figure 6E-I). Again, 10 out of 13 cells with lagging chromosomes were in telophase rather than in anaphase.

In conclusion, the analyses of the neocortex of the mice humanized for KIF18a and KNL1 and of the human organoids ancestralized for KIF18a and KNL1 show that the three modern human-specific amino acid substitutions in KIF18a and KNL1 prolong the mitotic metaphase of APs and decrease the occurrence of mitotic APs with chromosome segregation defects.

## Discussion

In the present study, we focus on modern human-specific amino acid substitutions in mitotic spindle and kinetochore proteins and their effects on the divisions of APs during neocortical development. There are several reasons for this. First, previous work has shown that metaphase is longer in modern human APs than in the apes(*15*). Second, this difference appears to be unique to the early stages of cerebral cortical tissue development, as it is not seen in non-neural cells nor in organoids at late stages(*15*). Third, among around 100 genes carrying essentially fixed amino acid substitutions unique to modern humans compared to archaic humans(*25*), the three genes studied here – *KNL1*, *KIF18a*, and *SPAG5* – are expressed, within the developing neocortex, predominantly in the ventricular zone, the primary germinal layer where APs reside(*40*). Fourth, the three proteins encoded by these genes carry a total of six amino acid substitutions unique to modern humans(*41*). This is more than expected given the small number of such amino acid differences. Fifth, these three proteins are involved in kinetochore function, a key process during metaphase(*22*).

When we introduce these modern human-specific six amino acid substitutions into mice, this results in a prolongation of metaphase in APs (Fig. 3J), reminiscent of what is seen when modern humans are compared to apes. The three substitutions in the two proteins KNL1 and KIF18a are enough to elicit the full effect, whereas the substitutions in either of the two proteins cause a metaphase prolongation of intermediate magnitude. In contrast, the three substitutions in SPAG5 have no effect. These data are consistent with the fact that the substitutions in KIF18a(*27*) and KNL1(*42, 43*) are located in, or close to, functionally relevant domains or motifs, whereas this is not the case for SPAG5(*34*) (see Figure 3A).

When the three modern human-specific amino acids in KNL1 and KIF18a are changed to the ancestral states shared by apes and archaic humans in human embryonic stem cells, this results in a shortening of the metaphase in APs in organoids, whereas other phases of AP mitosis are not affected (Fig. 4D). Thus, the humanization of *KNL1* and KIF18a in mice and the ancestralization of these genes in humans yield congruent results suggesting that the amino acid substitutions specific to modern humans cause a prolongation of the metaphase in APs. It is noteworthy that the magnitude of the shortening of AP metaphases seen in the ancestralized organoids is close to that previously observed between human and chimpanzee organoids (*15*). It is therefore possible that the three amino acid substitutions in KIF18a and KNL1 are largely responsible for the difference in AP metaphase length between present-day humans and apes. Given that the three ancestral amino acids are shared between chimpanzees and archaic humans, this implies that most, if not all, of the lengthening of AP metaphase evolved in modern humans subsequent to the divergence from archaic humans about half a million years ago.

A concern when performing genome editing is unintended effects that may affect the target sites, notably deletions(*39*) (Weisheit et al. 2020; Shin et al. 2017) that may not be detected when the target site is sequenced (Maricic et al., 2021). To exclude this, we determined the copy number of the locus after editing and verified that polymorphic positions up- and down- stream of each site remained heterozygous after editing. In preliminary experiments, we identified two cellular clones that carried the ancestralized sites but in one case in addition a deletion of one allele of *KNL1* and in the other case a deletion of one allele of *KIF18a*. Both these clones show a shortening of AP metaphase of similar magnitude as the diploid clones (Suppl. Fig. 6A). This suggests that mitotic APs harbor sufficient KIF18a and KNL1 protein so that one allele of either of the genes can be inactivated without detectable consequences for the aspects of cell division measured here. Corresponding decreases in SAC-positive kinetochores and increases in lagging chromosomes are also seen (Suppl. Fig. 6B-D).

Two other substitutions amino acid substitutions that have become essentially fixed in modern humans after the split from the ancestors of Neandertals and Denisovans have recently been analyzed. The first case is an amino acid substitution in the splice protein NOVA1 which is claimed to affect brain organoid morphology when edited to the ancestral state (Trujillo et al., 2021). However, this effect may be due to unintended hemizygous deletions (Maricic et al., 2021; but see also Herai et al. 2021). The second case is an amino acid substitution in the enzyme adenylosuccinate lyase, which decreases the stability of the protein and reduces purine biosynthesis when edited to the modern human-like state in mice and increases purine biosynthesis when ancestralized in human cells (Stepanova et al., 2021). It is unknown what further organismal effect this has but when humans and great apes are compared, the reduction in purine synthesis is particularly pronounced in the brain (Stepanova et al., 2021).

It is striking that the modern human-specific amino acid substitutions in KNL1 and KIF18a result in more kinetochores being associated with components of the SAC (Fig. 5; Suppl. Fig. 5) and in fewer lagging chromosomes or chromosomal fragments in mitotic APs (Fig. 6). Therefore, these substitutions are likely to result in higher accuracy of chromosome segregation in APs during neocortex development. This may have significant consequences. During the early stage of this development, APs undergo symmetric proliferative divisions that increase the AP pool size(*3–7, 44*). Any chromosome segregation error that occurs in an AP at this stage will therefore increase the number of APs with aberrant numbers of chromosomes. When APs switch to asymmetric divisions at the onset of cortical neurogenesis(*3–7, 44*), chromosomal aberrations may be transmitted to their progeny, *i.e.* the basal progenitors, cortical neurons and macroglial cells. Therefore, any radial unit derived from an AP with a chromosome segregation error, and hence the cortical column that contains this unit(*1, 45*), might be functionally affected. Mis-segregation may also result in apoptosis and loss of progenitors and the radial units they would have generated. In either case, consequences for the functionality of the entire neocortex area harboring an affected radial unit may ensue, for example for excitatory pyramidal neurons and their connections and projections. In addition, lingering chromosome segregation defects may have a variety of pleiotropic effects due to perturbed nuclear organization and imbalances in gene expression(*46, 47*). Such effects may tend to be more profound than the effects of somatic point mutations that may be involved, for example, in autism spectrum disorder(*48*). The present data imply that the probability of any such detrimental effects of chromosomal missegregation may be lower in modern humans than in Neandertals, Denisovans, and apes. Further work is needed to address if this is of importance for traits unique to modern humans.

## Materials and Methods

### Mice

All mouse experiments were performed according to the German Animal Welfare Legislation, (“Tierschutzgesetz”) after approval by the Federal State Authority Landesdirektion Sachsen. All procedures were overseen by the Institutional Animal Welfare Officer of MPI-CBG. Mice used for this study were kept in standardized and pathogen-free conditions at the Biomedical Services Facility (BMS) of MPI-CBG, with unlimited access to food and water, and in a 12 hours/12 hours light/dark cycle. Since the sex of embryos was unlikely to influence the results in this study, it was not taken into consideration. Embryonic day (E) 11.5 embryos were used throughout. E0.5 was defined as noon of the day of vaginal plug identification.

### Human tissue

Fetal human neocortex tissue (PCW 11-12) was obtained from the Klinik und Poliklinik für Frauenheilkunde und Geburtshilfe, Universitätsklinikum Carl Gustav Carus of the Technische Universität Dresden, with informed written maternal consent, and with approval of the Clinic’s Ethical Review Committees. PCW 11-12 corresponds to an early neurogenesis stage. The age of fetuses was assessed by ultrasound measurements of crown-rump length and other standard criteria of developmental stage determination. Due to protection of data privacy, the sex of the human fetuses from which neocortical tissue was obtained cannot be reported. The sex is unlikely to influence the results in the present study.

### Mouse genome engineering

For introduction of point mutations mimicking the human-specific changes, genome edited mouse lines were established using CRISPR-mediated homology directed repair (HDR) in isolated mouse zygotes, using the pronuclear injection (PNI) delivery method.

The following gRNAs with protospacers targeting the genomic positions of the desired changes in Kif18A, SPAG5 and KNL1 were either purchased as AltR crRNAs (IDT) or produced by *in vitro* transcription from a PCR template containing the T7 promoter followed by the gRNA sequence and the optimized gRNA scaffold(*49*) using the esiSCRIBE *In Vitro* Transcription Kit (Eupheria Biotech) following the manufacturer’s instructions:

Kif18a_R67K_sg1; CAAATTTTGATATTACTAAA

Knl1_H109R_sg1.1; CATTTGCATGTTTCCTTTCA

Knl1_G910S_sg2.1; TTTCTGAATGAACTTCTGTC

Spag5_P43S_sg1.2; GCACAGGTATGGGGTCAGCG

Spag5_E164G_sg2.2; AAGCTCTCTAAATGAATCTT

Spag5_D380H_sg3/cr3; AGGACAGTACTTCAGAGACA

As HDR templates, the following oligonucleotides were used:

Kif18a_R67K;

5’-

GCTTGGTTGTGTGTTCAAAAACTTCCATTTGAGTTGAAGTTTCATCAAAGACAGC

ATCAAATACAAACTTCAGATCTTTATTTTGTTTTTTAGTAATATCAAAATTTGTAG

TTTTCTTTCTGTGAAA-3’

Knl1_H109R;

5’-

TGCTGTCAGATCCATCTGGTTTTCATCTGAAAAGATGACTGTCTGATCATTTGCAT

GTTTGCGCTCGCGGTTGCAGTCTGTAATTGAGAACTATAATGAAAAAATAGAAATATTAGATCAAATATAAT-3’

Knl1_G910S; 5’-

ATTCTCTGGAAATGTAATACTCTTGTTTTTGAGGCTAAGGCTTTTTCTTCTCTGAGTTTTGGATGACAGAAGTTCATTCAGAAATCCAGGCTTTTGTGCATCATTAGTTTCCTCTCCAAGCCC-3’

Spag5_P43S;

5’-

AGTACTCCTCTCCGAGAGCTTAAACTACAGCCCGAGGCCCTCGCCGATTCAGGGAAAGGTAGTAGCATGATTAGTGCACTGACCCCATACCTGTGCAGGCTGGAGCTGAAGGTGAGATTTTTAGCTTCACAACTGGGCT-3’

Spag5_E164G;

5’-

TGTTAGGCTGTCTTCCACACAAGGTGCCACCTCCTTCCCCACCAGATCTTCTAGCCCCAAGCTGCCATTTAGAGAGCTTTTAGTTCTCTCTGCTGTGGTATCTAAGTGAGCCTGGAGCATCAGGTC-3’

Spag5_D380H;

5’-

CAGGATGCTGCCGTTGGCAACACACCCCTCGCCACGTGTTCTGTGGGCACTTCCTTTACTCCTCCAGCACCACTGGAGGTAGGCACAAAACACAGTACTTCAGAGACAGAGCGCCTCCTCTTGGGGT-3’

Donor / Background / CRISPR/Cas9-Injection. C57Bl6/NCrl mice were mated after superovulation (46 hours between PMSG and HCG) at the midpoint of the dark period (12 hours/12 hours, 4 am to 4 pm light circle). After positive plug detection in the morning, the cumulus complexes were isolated and zygotes removed with a treatment of hyaluronidase (final concentration of 0.1% (801 units/ml)). CRSIPR/Cas9 solutions were injected into the male pronucleus of fertilized zygotes by using a motor-driven manipulator-based microinjection stage. About two hours after injections, the surviving embryos were transferred into Crl:CD1(ICR) pseudopregnant recipient female mice (ca. 20 embryos per recipient).

Embryo Transfer. The recipient mice were mated with vasectomized males Crl:CD1 (ICR). After detection of a copulation plug in the morning of the transfer day, pseudopregnant mice were used for unilateral surgical embryo transfer into the oviduct as described(*50*). Anesthesia was induced by intraperitoneal injection of Ketamine/Xylacine and Acepromazine (*51*).

Mouse lines for single genes and gene combinations. hKNL1–hKIF18a as well as hSPAG5 mice, each carrying three missense changes, were generated by first combining guide RNAs for all three changes, and then repeating cycles for those changes not obtained in the previous rounds. The hKNL1 and hKIF18a lines were generated independently from the combined hKNL1–hKIF18a line, as *Knl1* and *Kif18a* reside on the same mouse chromosome. The hKNL1–hKIF18a–hSPAG5 line was generated by crossing the hKNL1–hKIF18a and hSPAG5 lines.

Candidate mice for each line were genotyped with primers and conditions described in Supplementary Table 1. Positive mice for each line were back-crossed with wild type C57Bl6/NCrl mice and genotyped for at least six generations before experiments.

#### Genotyping of mice

Genomic DNA from tail biopsies and buccal swabs of three-week-old pups were prepared using the Sigma Extract-N-Amp tissue kit following the manufacturer’s instruction but using only half of the described volumes. The target regions were amplified by PCR in a total volume of 25 µl using the Phusion Flash Mastermix (Thermo Scientific) with 2 µl genomic tail DNA, 0,25% DMSO and 0,5 µM forward and reverse primer. Primer sequences (target-seq-primer) and PCR conditions are in Supplementary Table 1. Products were Sanger sequenced either with the forward and reverse PCR primers or if required with internal sequencing primers. Primary candidates have been identified by sequence alignments of potentially modified PCR fragments against the wild type and modified reference sequence. The products from putatively edited mice were cloned (TOPO) and verified by Sanger sequencing of 10 to 20 randomly selected clones. Founders carrying the respective genome modification either homo- or heterozygously were propagated and genotyped by Sanger sequencing of PCR fragments. From the second generation onwards, allele-specific PCRs were performed on 2 µl crude buccal DNA in a total of 10 µl using the REDExtract-N- Amp^TM^ PCR ready Mix^TM^ (Sigma) according to the manufacturer’s protocol using 0.5 µM forward, reverse, and mutant primer. The readout is done on standard agarose gels. Primers and PCR conditions are in Supplementary Table 1.

#### Expression analysis of edited genes in humanized mice

For Sanger sequencing and quantitative real-time PCR (qPCR), total RNA was isolated from E11.5 wildtype and humanized mouse dorsolateral telencephalon using the RNAeasy Micro Plus Kit (Qiagen) according to the manufacturer’s instructions. cDNA was synthesized using the Maxima first-strand cDNA synthesis kit (Thermo Scientific). The expression of *Kif18a*, *Knl1* and *Spag5* in the wildtype and humanized embryonic mouse telencephalon was confirmed by PCR, using generated cDNA as templates, followed by Sanger sequencing of the PCR products. Wildtype and gene-edited sequences were mapped to the *Kif18a*, *Knl1* and *Spag5* genes, respectively, using Geneious Prime (version 2020.2.4). Data showed the expression of only the wildtype *Kif18a*, *Knl1* and *Spag5* in E11.5 wildtype embryonic mouse telencephalon, and the expression of only the gene-edited *hKif18a*, *hKnl1* and *hSpag5* in E11.5 humanized hKIF18a- hKNL1-hSPAG5 (see Figure 3A) embryonic mouse telencephalon (Supplementary Figure 2A-F). Relative mRNA levels of *Kif18a*, *Knl1* and *Spag5* to *Actb* in wildtype and humanized embryonic mouse telencephalon were quantified by qPCR, which was performed using the FastStart essential DNA green master (Roche) and LightCycler 96 Instrument (Roche). Gene expression analyses by qPCR showed that the relative mRNA levels of *Kif18a*, *Knl1* and *Spag5* were not significantly different between E11.5 wildtype and humanized embryonic mouse telencephalon (Supplementary Figure 2G-I).

The following primers were used for PCR:

Forward primer for Kif18a (5’-AGAAAAGGCGGTGCAGTTCT-3’),

Reverse primer for Kif18a (5’- GGAATCTTCCCGGACAGCAA-3’);

Forward primer for Knl1 H159R (5’-CCCCAGACAAGTCAAGCAGAA-3’),

Reverse primer for Knl1 H159R (5’-TCAACTCCATACACTCATTGCC-3’);

Forward primer for Knl1 G1086S (5’-TGGATATCACCAAGAGTTGCAC-3’),

Reverse primer for Knl1 G1086S (5’-CAAAACTGAAGCCCTTTCTGTC-3’);

Forward primer for Spag5 D410H and E162G (5’-TGAATCTCGGTTTGTCGCCT-3’),

Reverse primer for Spag5 D410H and E162G (5’-TTCCTCCCCTGGATCGACAT -3’);

Forward primer for Spag5 P43S (5’-GTTCAAATAGAGGCGGCGGG-3’),

Reverse primer for Spag5 P43S (5’-GTTCTTTCCCACCAGCTACAAG-3’).

The following primers were used for qPCR:

Forward primer for Kif18a (5’-CAAACTCAGGACCACTTGCTGT-3’),

Reverse primer for Kif18a (5’-ATGGGAACGAGAAGACACTGC-3’);

Forward primer for Knl1 (5’-CCTCTGGGGGAGATGGCTACAT-3’),

Reverse primer for Knl1 (5’-GATGGACTTTGTTGGGCTGAGA-3’);

Forward primer for Spag5 (5’-GTCTCACCCTCTTCTTACAGGC-3’),

Reverse primer for Spat5 (5’-GCTGGTTCTGGCACTTCATCTA-3’);

Forward primer for Actb (5’-CGGGACCTGACAGACTACCTC-3’),

Reverse primer for Actb (5’-GGTGGTGAAGCTGTAGCCACG-3’).

### Genome engineering of human H9 cells

We used an iCRISPR-Cas9n line carrying a doxycycline inducible Cas9D10A nickase in the *AAVS1* locus from H9 human embryonic stem cell line (female, WiCell Research Institute, ethics permit AZ 3.04.02/0118) which we generated as previously described (*52, 53*) and that carries a K3753R (KR) mutation in the *PRKDC* gene to increase editing efficiency as described (*54*). Cells were grown on Matrigel (Corning, 35248) in mTeSR1 medium (StemCell Technologies, 05852) supplemented with 2 µg/ml doxycycline (Clontech, 631311) for 3 days prior to electroporation and to induce Cas9 nickase expression. Electroporation of oligonucleotides and, in cases when editing with the nickase did not work, ribonucleoprotein (RNP) was carried out using 1 million cells, 78 pmol electroporation enhancer (IDT), 160 pmol of the gRNA (crRNA/tracR duplex for Cas9 and crRNA for Cpf1), 200 pmol of the respective single-strand DNA (ssDNA) donor, and 252 pmol Cas9-HiFi or Cpf1-Ultra (both IDT) using the B-16 program of the Nucleofector 2b Device (Lonza) in cuvettes for 100 µl Human Stem Cell nucleofection buffer (Lonza, VVPH- 5022). Cell were counted with Countess automated cell counter (Invitrogen).

*KIF18A* was ancestralized using iCRIPSR-Cas9 nickase with two gRNAs and ssDNA donor. The ancestralization of *KNL1*_G1086S was done by Cpf1-Ultra RNP and of KNL1_H109R by Cas9-HiFi RNP. The guides and the donors were purchased as Alt-R crRNAs and DNA oligonucleotides (IDT):

KIF18A_sg1: 5’-TGTTATAAAGAAACAAAATA

KIF18A_sg2: 5’-GATTTGTAGTTTTCTTTCCA

KNL1_H109R_sg1: 5’-ATTTGCATGTTTCCTTTCAC

KNL1_G1086S_sg1: 5’-TGAATGAACCTCTATCAAGCA

ssDNA KIF18A_K67R: 5’-

AGTTGACGTTTCATCAAAAACAGCATCAAATACAAATTTAAGATCCTTGTTTTGTCTCTTTATAACATTTTGATTTGTAGTTTTCTTTCCATGGAAAAAACTGACT

ssDNA KNL1_H159R: 5’-

TGTTGTAAAATGCCTTTTTAAAAGTTTGCTTTTGTTCTGATCTCTTATATTTTGCTTTCATTATAGTTTTCAATTATAGAACATACCCATGAAAGGAAACATGCAAATGACCAGACAGTCATTTTTT

ssDNA KNL1_G1086S:

5’-

TCTTGTCATTTTTTAGCTTAAGGCTTTTTCTTCTCTGACTTTTGCCTGATAGAGGTTCGTTCAGAAATCCAGGACTTTGTACATCTTTGA

To derive colonies from single cells, cells were incubated with StemFlex with supplement (Gibco, A3349401) and CloneR (StemCell Technologies, 05888) for one day and then sorted with a Cytena cell printer.

Validation of edited cell lines. Genomic DNA of each colony was isolated, a region of ∼200 bp around the cut site was amplified and sequenced, and from sequences the editing state was evaluated as described (*54*).

KNL1_G1086S_F: 5’-

ACACTCTTTCCCTACACGACGCTCTTCCGATCTTGATTCAAGCAAACCAACGTGT

KNL1_G1086S_R: 5’-

GTGACTGGAGTTCAGACGTGTGCTCTTCCGATCTTGATGTCAGGTCCATCTGGT

KNL1_H159R_F: 5’-

ACACTCTTTCCCTACACGACGCTCTTCCGATCTTCCCAAATTGAAACAAATACCTG

KNL1_H159R_R: 5’-

GTGACTGGAGTTCAGACGTGTGCTCTTCCGATCTTTTTTGATCCCAAACAAGAAG AA

KIF18A_F: 5’-

ACACTCTTTCCCTACACGACGCTCTTCCGATCTTCAGCCACTTGCAAAAACATCA

KIF18A_R: 5’-

GTGACTGGAGTTCAGACGTGTGCTCTTCCGATCTTAGAAGCCCCTGCCAAATGG

Deletions bigger than ∼200bp and therefore not detected by the amplicon sequencing(*39, 55*) were identified by digital PCR using primer pairs and probes that anneal within the sequenced target regions. A primer pair and probe were also designed for the gene *FOXP2* to serve as a diploid reference. The master mix for droplet digital PCR (ddPCR) amplification included 1× ddPCR Supermix for probes (no dUTP, Bio-Rad), 0.2 1M of each primer and 0.2 1M probe (both IDT) for target and reference, together with 1 1l genomic DNA in QuickExtract DNA Extraction Solution (Lucigen). After droplet generation with the QX200 Droplet generator (Biorad), the PCR reaction was for 5 min at 95°C, followed by 42 cycles of 35 s at 95°C (at a ramp rate of 1.5°C/s) and 65 s at 54°C for *KIF18A* and *KNL1*_G1086S or 65 s at 56°C for *KNL1*_H109R (at a ramp rate of 1.5°C/s), and 5 min at 98°C. Readout was in a QX200

Droplet reader (BioRad) and allele copy numbers were determined relative to the *FOXP2*

reference and unedited controls.

KNL1_G1086S_ddPCR_L: 5’-AAGATGTACAAAGTCCTGGAT

KNL1_G1086S_ddPCR_R: 5’-CTCTTTATCCTCCAGGGC

KNL1_G1086S_probe1: 6-FAM 5’-TGGATATTACCCAGAGTTGTATGGTGGA BHQ_1

KNL1_H159R_ddPCR_L: 5’-TATGAAACCATCACTGAGGA

KNL1_H159R_ddPCR_R: 5’-CTGAAAAAATGACTGTCTGGTCA

KNL1_H159R_probe1: 6-FAM 5’-TGCCTTTTTAAAAGTTTGCTTTTGTTCTGA BHQ_1

KIF18A_ddPCR_L: 5’-ATATCCATTCAAAAAACTACGAA

KIF18A_ddPCR_R: 5’-CCATGGAAAGAAAACTACAAA

KIF18A_ddPCR_probe1: 6-FAM 5’-TGAGTTGACGTTTCATCAAAAACAGCATCA BHQ_1

FOXP2_ddPCR_F: 5’-GCAACAGCAATTGGCAGC

FOXP2_ddPCR_R: 5’-CAGCGATTGGACAGGAAGTG

FOXP2_ddPCR_probe1: HEX 5’-AGCAGCAGCAGCATCTGCTCAGCCT BHQ_1

To exclude colonies that experienced a loss-of-heterozygosity at the region near the CRISPR cut site (*39*), heterozygous variable positions upstream and downstream of the respective target site were amplified and sequenced.

**KNL1_G1086S** (upSNP distance: 809 bp and 868 bp; downSNP distance: 254 bp)

KNL1_G10862_upSNP_F: 5’-

ACACTCTTTCCCTACACGACGCTCTTCCGATCTAAGCAATCCCACACCTGACT

KNL1_G10862_upSNP_R: 5’-

GTGACTGGAGTTCAGACGTGTGCTCTTCCGATCTCTTTCGTCCTCTAACACATCCA

KNL1_G10862_downSNP_F: 5’-

ACACTCTTTCCCTACACGACGCTCTTCCGATCTGCCCTGGAGGATAAAGAGGA

KNL1_G10862_downSNP_R: 5’-

GTGACTGGAGTTCAGACGTGTGCTCTTCCGATCTGAATCGGTGACTTCCAGATCA

**KNL1_H159R** (upSNP distance: 2153 bp and 2192 bp; downSNP distance: 1317 bp)

KNL1_H159R_upSNP_F: 5’-

ACACTCTTTCCCTACACGACGCTCTTCCGATCTAAGACCCTTCAGAATCCTACCC

KNL1_H159R_upSNP_R: 5’-

GTGACTGGAGTTCAGACGTGTGCTCTTCCGATCTTGTATCCTCTGTGTGGCTGAA

KNL1_H159R_downSNP_F: 5’-

ACACTCTTTCCCTACACGACGCTCTTCCGATCTGCAGAGCCTGTCAAATCCTT

KNL1_H159R_downSNP_R: 5’-

GTGACTGGAGTTCAGACGTGTGCTCTTCCGATCTCCAGGTGTCGGTTAAGCTGT

**KIF18A** (upSNP distance: 24343 bp; downSNP distance: 30555 bp)

KIF18A_upSNP_F: 5’-

ACACTCTTTCCCTACACGACGCTCTTCCGATCTGTGGGTAAGGGAGTGGGAAT

KIF18A_upSNP_R: 5’-

GTGACTGGAGTTCAGACGTGTGCTCTTCCGATCTGCTTCAGAGCCTGCACTCTT

KIF18A_downSNP_F: 5’-

ACACTCTTTCCCTACACGACGCTCTTCCGATCTGCAACCGTTTTAGCCAAGAT

KIF18A_downSNP_R: 5’-

GTGACTGGAGTTCAGACGTGTGCTCTTCCGATCTCTCCCCGAATGTCTTTCTCA

To find large-scale chromosomal deletions, duplications or karyotype abnormalities we performed “shallow” whole genome sequencing to an average genome coverage 0.10-0.15 of edited and non-edited colonies as well as the cell line from which the colonies were derived, as described (*54*).

Expression of *KIF18a* and *KNL1* in edited and non-edited cell lines as well as their “mother” cell line was measured with CellsDirect™ One-Step qRT-PCR Kit (Invitrogen) using TaqMan probes for *KIF18a* (FAM Hs0105428_m1), *KNL1* (FAM Hs00538241_m1) and a “housekeeping gene” *PPIB* (VIC Hs00168719_m1). The producer’s protocol was followed to set up reactions. For a subset of reactions reverse transcriptase was inactivated by 15 min-incubation at 70°C. The cycler program for RT-qPCR was: 50°C for 10 min, 95°C for 2 min, then 50 cycles of: 95°C for 3 s, 53°C for 15 s and 60°C for 30 s.

### Generation of cerebral organoids

Organoids were generated from the H9 ESCs and cultured as described (*56, 57*) with the exceptions that TrypLE was used instead of accutase, ROCK inhibitor was used on the first two days, mTeSR1 was used instead of low-bFGF hESC medium in 96-well ultra-low attachment plates during embryoid body (EB) formation, neural induction medium was given from day 4, embedding was on day 9 with B27 in the differentiation medium, and vitamin A was added on day 15. On day 27-32 they were fixed, embedded, cryosectioned and prepared for immunohistofluorescence as described(*15*). Fixation was with 4% PFA in 120 mM phosphate buffer pH 7.4 for 2 h at room temperature followed by overnight incubation at 4°C.

### Neocortex tissue

E11.5 mouse neocortex tissue was dissected at room temperature in PBS. The dorsolateral telencephalon, at a medial position along the rostro-caudal axis, was used for live tissue imaging (unfixed) and immunohistofluorescence (fixed). Mouse neocortex was fixed, embedded, cryosectioned and prepared for immunohistofluorescence as described(*15, 58*), except that fixation was with 1% PFA in 120 mM phosphate buffer pH 7.4 for 30 min at room temperature. Human neocortex tissue was fixed with 4% PFA in 120 mM phosphate buffer pH 7.4 for 2 h at room temperature followed by overnight incubation at 4°C.

### Immunohistofluorescence

Immunohistofluorescence was performed as described(*15, 58*). Analyses of SAC-positive kinetochores, as well as of lagging chromosomes, was performed using 3D stacks of serial confocal sections, of typically 512 × 512 pixels × 16–25 z-sections (xyz sampling: 0.09 × 0.09 × 0.75 µm). The following primary antibodies were used: anti-BUBR1 mouse monoclonal (1:100 dilution; BD Biosciences), CREST anti-centromere (kinetochore) human polyclonal (1:50; Antibodies Incorporated), anti-MAD1 mouse monoclonal (1:100; Millipore), anti-FOXG1 rabbit polyclonal (1:300; Abcam). The secondary antibodies used (1:500 dilution) in combination with DAPI staining were all donkey-derived, conjugated with Alexa 488, 555 or 647 (Life Technologies), except goat anti-human conjugated with Alexa 488 (Life Technologies).

### Live imaging

Live tissue and cell imaging were performed as described(*15, 58*). In short, freshly dissected developing neocortex tissue or organoids were embedded in agarose (Sigma, Germany), sectioned (∼200 mm) with a vibratome (Leica, Germany), embedded in type Ia collagen (Cellmatrix, Japan), mounted in glass bottom microwell dishes (MatTek, Germany), and incubated with Hoechst 33342 (Sigma) as vital DNA dye. Tissue slices in the dish were cultured in a microscope stage incubation chamber (Pecon, Germany) at 37°C. The H9 ESC monolayer cultures were mounted in glass bottom microwell dishes coated for 1h with

Matrigel (BD Biosciecne), and imaged under standard cell culture conditions. Analysis of mitotic phase durations were performed on time-lapse 3D stacks of serial 2-photon sections of typically 512 × 512 pixels × 7-9 z-sections (xyzt sampling: 0.7 × 0.7 × 2.2 µm × 1.5–2.5 min), acquired for 6–12 h.

### Microscopy

Fixed – as well as live – images were recorded with a Zeiss LSM 780 NLO single/multi-photon point scanning system, with tunable pulsed near-infra-red laser (Chameleon Vision II) for multiphoton excitation, and using 63× Plan-Apochromat 1.4 N.A. oil or 40× C-Apochromat 1.2 N.A. W objectives (Carl Zeiss, Germany).

### Quantifications

Images were viewed and prepared with ImageJ (http://imagej.nih.gov/ij/). Brightness and contrast were recorded and adjusted linearly. Measurements of mitotic phases were performed as described (*15*). In short, the time of chromosome congression toward a metaphase plate is referred to as “congression” or “prometaphase”; the time between the formation of a tight metaphase plate and chromatid segregation onset is referred to as “metaphase”; the time between the onset of chromatid segregation and the onset of general chromosome decondensation onset is referred to as “anaphase”; the time between the beginning of chromosome decondensation until a level indistinguishable from interphase is referred to as “telophase”. Given the low incidence of lagging chromosomes, the percentage of AP divisions with lagging chromosomes was determined by adding all cells from all independent experiments per tissue to a single, cumulative percentage for each.

Quantifications of AP mitotic phases, SAC markers and lagging chromosomes in mouse neocortex were performed by a blinded observer (F.M.B.), and lagging chromosomes in mouse neocortex and organoids were cross-checked by another blinded observer (C.H.).

Quantifications of AP metaphase durations in organoids were reproduced by a blinded observer (W.B.H).

### Statistical analysis

Cell and tissue data were plotted and analyzed with GraphPad Prism (La Jolla, CA). Since at least some data sets in most experiments did not pass normality tests (Shapiro-Wilk and Kolmogorov-Smirnov tests), non-parametric Mann–Whitney U-tests were used for two groups of observations and Kruskal–Wallis tests with Dunn’s correction for multiple comparisons for three or more groups. For gene-expression analysis in mice the two-tailed unpaired Student’s t-test was used.

## Acknowledgments

We thank members of the Huttner and Pääbo groups for useful discussions. We apologize to authors whose work we could not refer to due to space constrains. We thank the Services and Facilities of the Max Planck Institute of Molecular Cell Biology and Genetics for expert technical support, notably J. Peychl and his team of the Light Microscopy Facility and J. Helppi and his team of the Biomedical Services (BMS), and the teams in the Transgenics, Genome Engineering and Genotyping Facilities. We thank Damian Wollny for generating the H9 iCRISPR cell line.

## Author contributions

F.M-B., conceived, designed, performed and analyzed experiments, designed gene-edited mice, optimized and generated cerebral organoids, wrote the manuscript; P.K. and D.M. did genome engineering of human stem cells and validated edited cellular clones; J.P., optimized and generated cerebral organoids; R.N., designed and generated gene-edited mice; M.S., designed gene-edited mice; S.W., designed mouse genotyping; C.E.O., generated cerebral organoids; C.H. cross-checked data; L.X., analyzed gene expression in mice; P.W. provided fetal human tissue and information relevant for data interpretation; S.R. did genome engineering of human stem cells and validated edited cellular clones; T.M., validated edited cellular clones; W.B.H supervised the study, cross-checked and reproduced data, wrote the manuscript; S.P. supervised the study and wrote the manuscript.

## Funding

This work was funded by the Max Planck Society (W.B.H, S.P.), ERA-NET NEURON (MicroKin) (W.B.H.) and the NOMIS Foundation (S.P.).

## Competing interests

The authors declare that they have no competing interests.

## Supplementary Information

### Supplementary Figures and Figure Legends

**Supplementary Figure 1.**
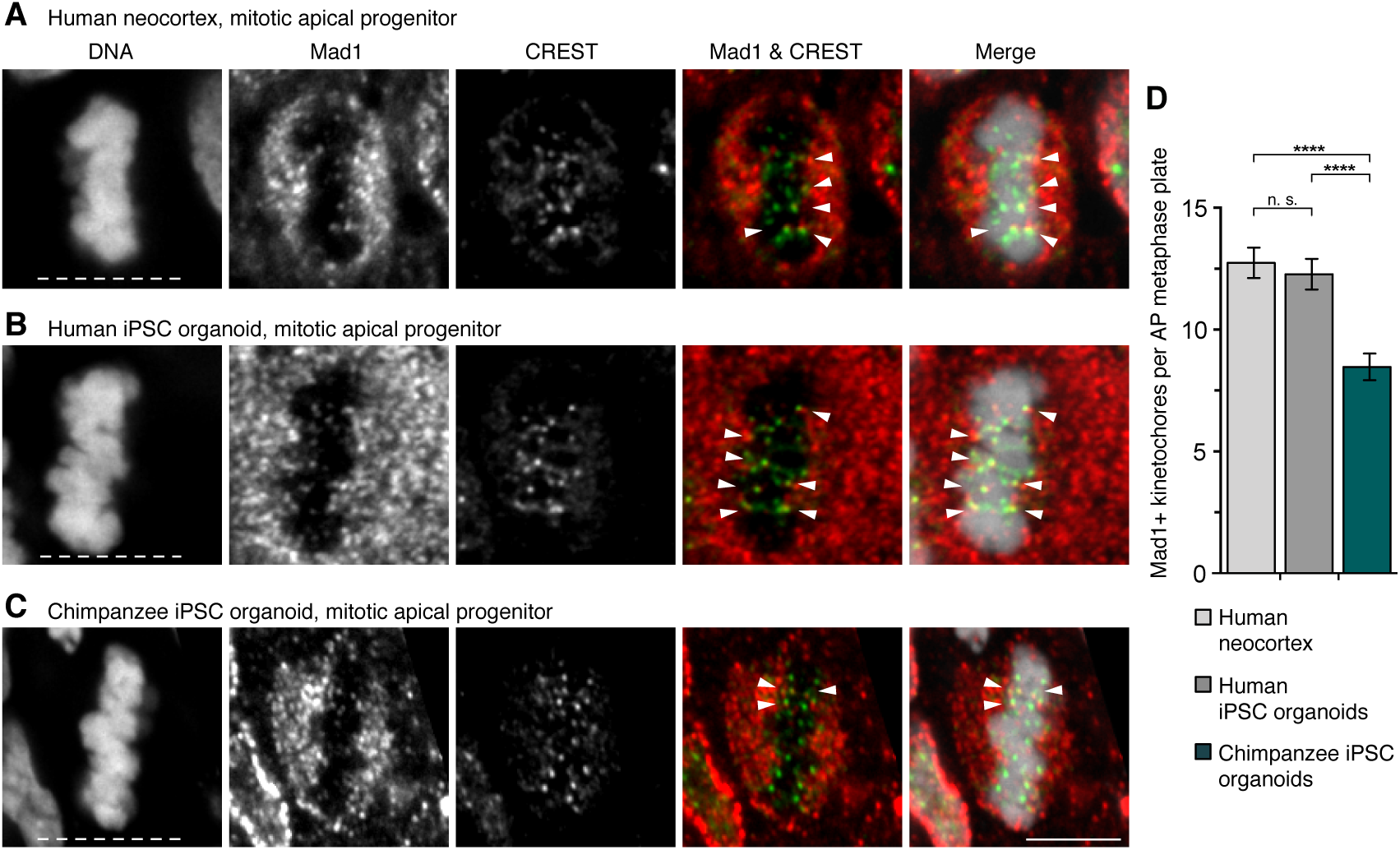
SAC-positive kinetochores in modern human and chimpanzee APs. (**A–C**) Mitotic APs in metaphase stained with DAPI and immunostained for the SAC marker Mad1 (red in merges) and the kinetochore marker CREST (green in merges). Arrowheads, overlap of Mad1 and CREST immunoreactivity within the metaphase plate. (**A**) GW11 modern human neocortex; (**B**) day 30 modern human iPSC-derived cerebral organoid; (**C**) day 31 chimpanzee iPSC-derived cerebral organoid. White dashed lines, ventricular surface. Scale bar, 5 μm. (**D**) Quantification of Mad1-positive kinetochores per AP metaphase plate for the tissues in A–C (GW11-12, days 30-32). Data are the mean ± SEM of ≥37 APs from three independent experiments with three neocortex samples, and ≥50 APs from ≥3 independent experiments with ≥5 organoids per species. Brackets with ****, p <0.0001; n. s., non-significant (Kruskal-Wallis test with Dunn’s multiple comparisons correction).

**Supplementary Figure 2.**
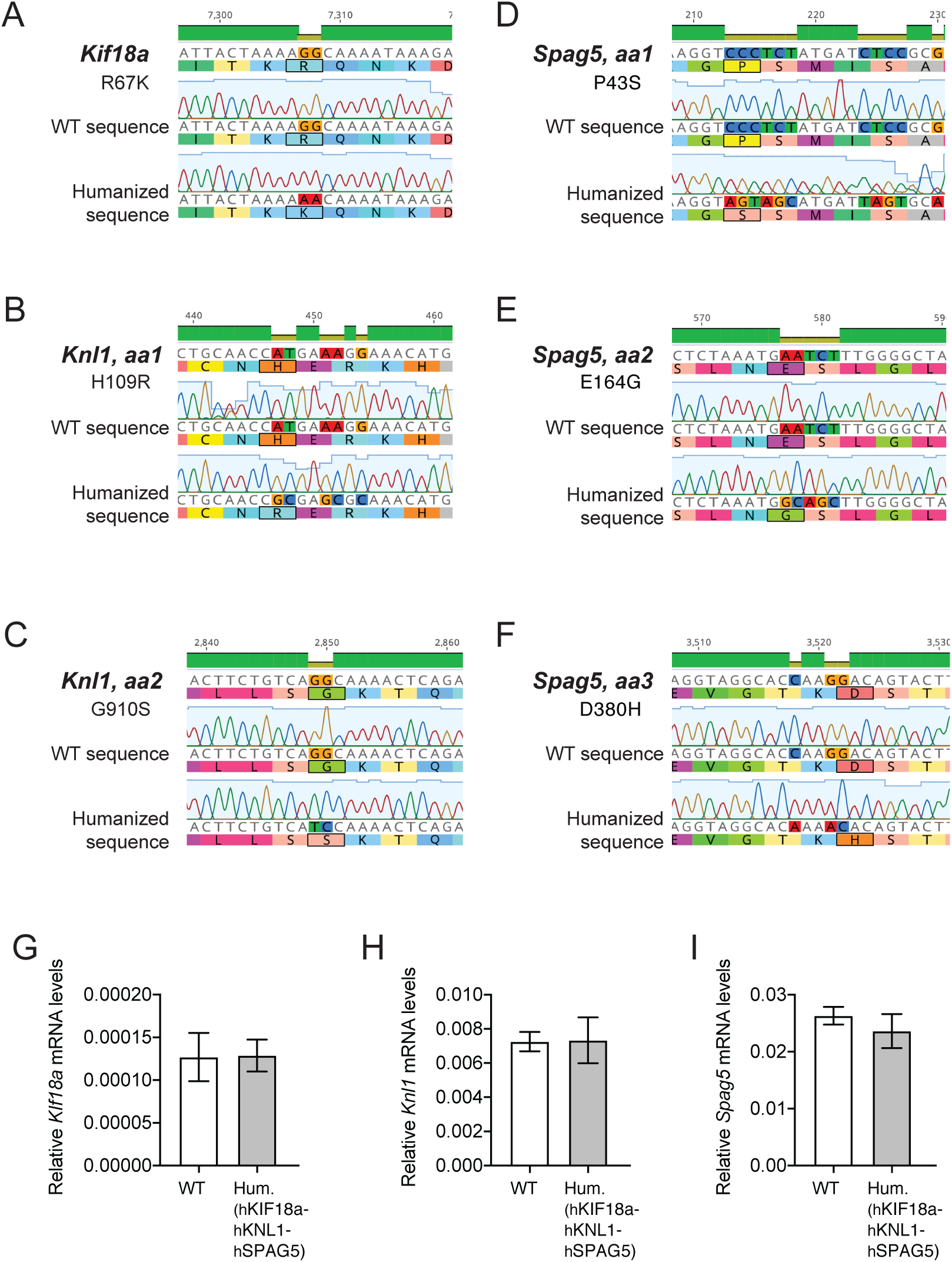
*Kif18a*, *Knl1* and *Spag5* mRNA expression in wild-type and humanized E11.5 mouse telencephalon. (**A**) *Kif18a* sequences in wildtype (WT) and humanized (h or Hum, hKif18a-hKnl1-hSpag5 (see also Figure 3A) E11.5 mouse telencephalon, aligned with the mouse *Kif18a* gene. (**B**, **C**) *Knl1* sequences for the amino acid (aa) positions 1 and 2 (aa1 & aa2) in WT and humanized 11.5 mouse telencephalon, aligned with the mouse *Knl1* gene. (**D-F**) *Spag5* sequences for the aa positions 1, 2 and 3 (aa1, aa2 & aa3) in WT and humanized E11.5 mouse telencephalon, aligned with the mouse *Spag5* gene. (**G-I**) qPCR analysis of the expression of *Kif18a* (G), *Knl1* (H) and *Spag5* (I), relative to the expression of *Actb,* in the WT and humanized E11.5 mouse telencephalon. Data are the mean of four biological replicates, each of which contains one or two telencephala. Error bars indicate SD, two-tailed unpaired Student’s t-test. p = 0.9210 (G), p = 0.9207 (H), p= 0.1603 (I).

**Supplementary Figure 3.**
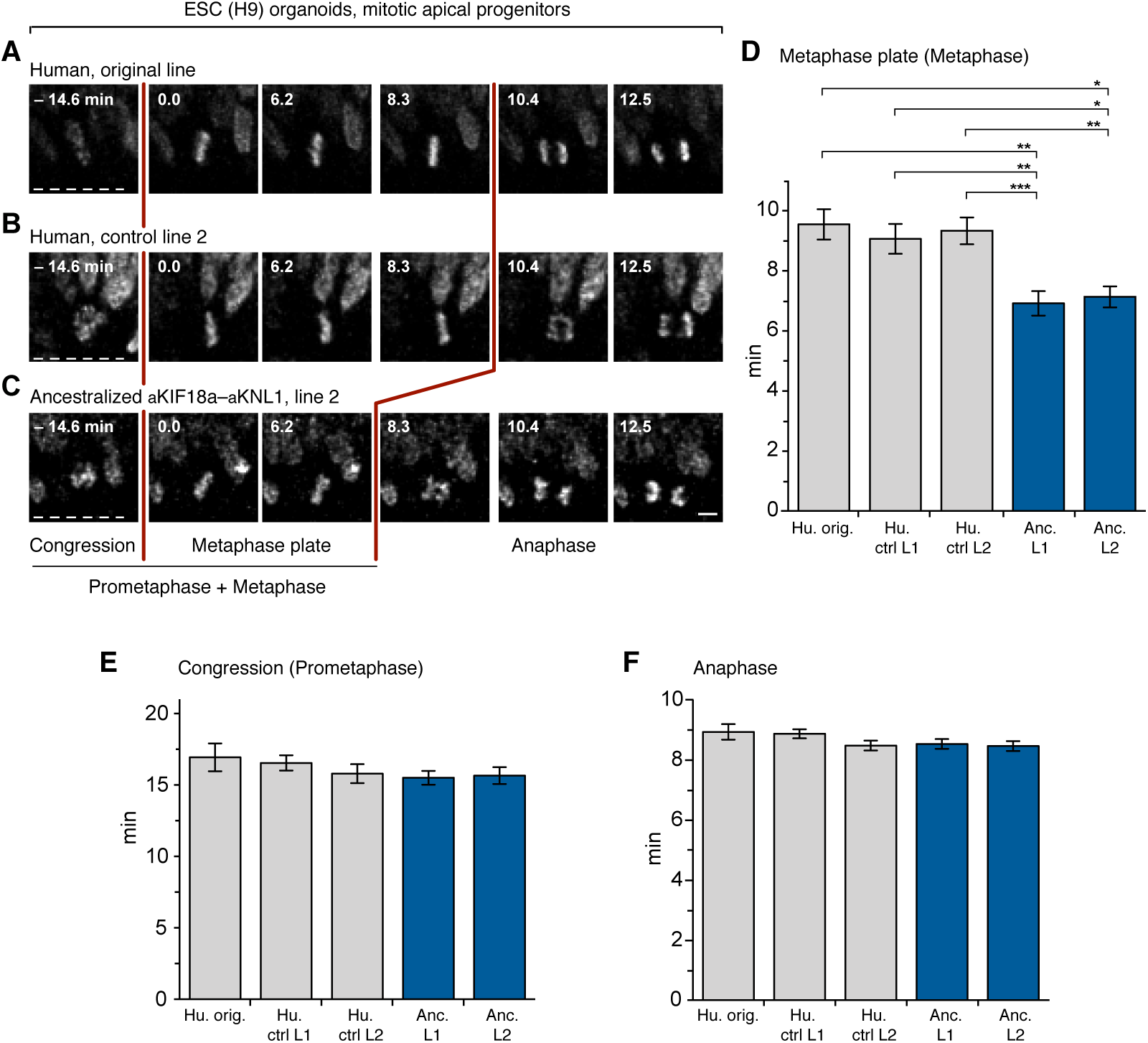
AP mitosis in organoids ancestralized for KIF18a and KNL1. (**A-C**) Live-tissue imaging of the indicated mitotic phases of apical progenitors in organotypic slice cultures of day 27-30 cerebral organoids grown from the indicated ESC lines. (**A**) original, non-edited human H9 line (Mod. hu. orig. in D-F); (**B**) the control line 2 (line 1 is shown in Figure 4B); (**C**) the ancestralized aKIF18a–aKNL1 line 2 (line 1 is shown in Figure 4C). Zero (0) min is metaphase plate onset. Time-lapse intervals are 2.08 min. Red lines indicate the duration of metaphase. White dashed lines, ventricular surface. Scale bar, 5 μm. Times (**D**) between metaphase plate onset and chromatid segregation onset (referred to as “Metaphase plate” or “Metaphase”); (**E**) between chromosome congression onset and metaphase plate onset (referred to as “Congression” or “Prometaphase”); and (**F**) between chromatid segregation onset and chromosome decondensation onset (referred to as “Anaphase”), for apical progenitors in organoids grown from each of the indicated cell lines (crtl, control; L, line, Anc., ancestralized). Data are the mean ± SEM of ≥52 APs from ≥2 independent experiments, with a total of ≥4 organoids, for each of the five lines. Brackets with *, p <0.05; ** p <0.01; ***, p <0.001 (Kruskal-Wallis test with Dunn’s multiple comparisons correction).

**Supplementary Figure 4.**
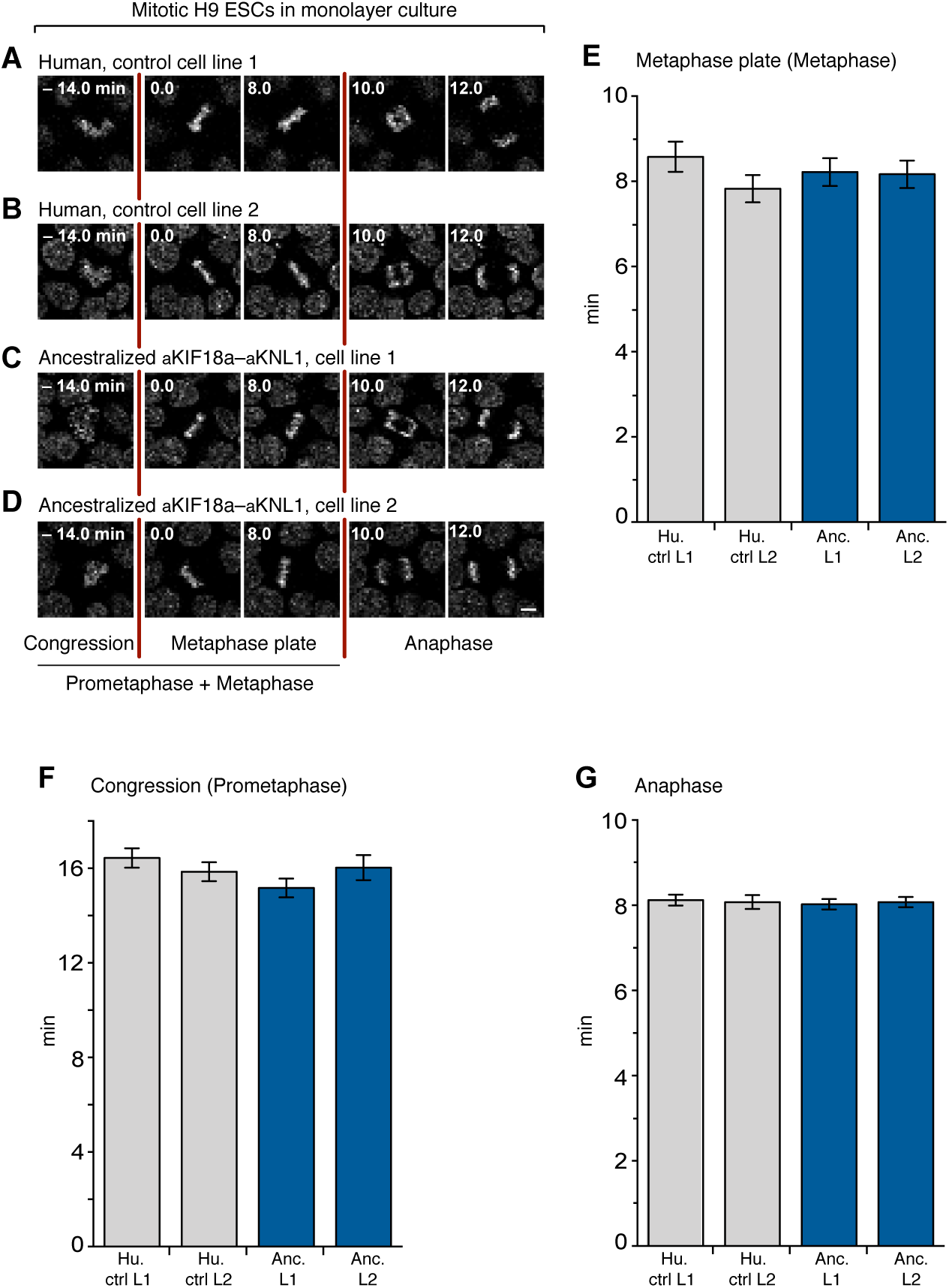
Mitosis of human ESCs, ancestralized for KIF18a and KNL1, in monolayer culture. (**A-D**) Live-tissue imaging of the indicated mitotic phases of the indicated H9 ESCs in monolayer culture. (**A**) Control ESC line 1; (**B**) control ESC line 2; (**C**) ancestralized aKIF18a–aKNL1 ESC line 1; (D) ancestralized aKIF18a–aKNL1 ESC line 2. Zero (0) min is metaphase plate onset, time-lapse intervals are 2 min. Red lines indicate the duration of metaphase. Scale bar, 5 μm. Times (**E**) between metaphase plate onset and chromatid segregation onset (referred to as “Metaphase plate” or “Metaphase”); (**F**) between chromosome congression onset and metaphase plate onset (referred to as “Congression” or “Prometaphase”); and (**G**) between chromatid segregation onset and chromosome decondensation onset (referred to as “Anaphase”), for each of the four H9 ESC lines described in (A–D; Hu., human; Anc., ancestralized; ctrl, control, L, line). Data are the mean ± SEM of ≥81 cells from three independent experiments, for each of the four lines.

**Supplementary Figure 5.**
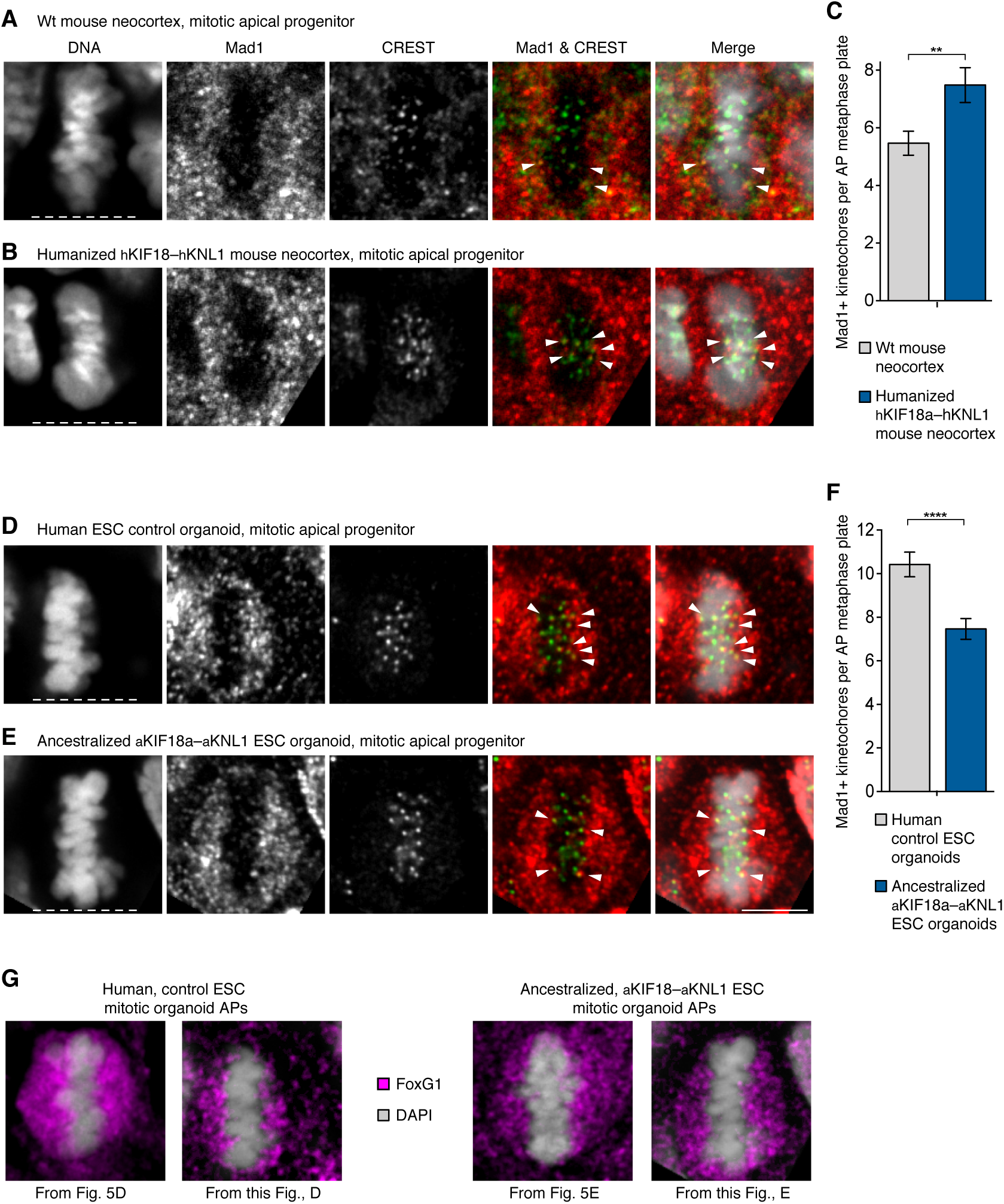
SAC-positive kinetochores in mice humanized for Kif18a and Knl1 and organoids ancestralized for KIF18a and KNL1. (**A, B, D, E**) Mitotic APs in metaphase stained with DAPI and immunostained for the SAC marker Mad1 (red in merges) and the kinetochore marker CREST (green in merges). Arrowheads indicate overlap of BubR1 and CREST immunoreactivity within the metaphase plate. (**A**) Neocortex of E11.5 wild type (wt, ancestral-like) mouse; (**B**) neocortex of E11.5 mouse humanized for *Kif18a* and *Knl1*; (**D**) modern human non-edited day 28 cerebral organoid (control line 2); (**E**) day 29 organoid ancestralized for KIF18a and KNL1 (edited line 2). White dashed lines, ventricular surface. Scale bar, 5 μm. (**C**) Number of Mad1-positive kinetochores per AP metaphase plate in mouse neocortex as described in (A, B). Data are the mean ± SEM of ≥32 APs from three independent experiments, with a total of ≥5 neocortices, for each of the two types of mice. Bracket with **, p <0.01 (Mann-Whitney U test). (**F**) Number of Mad1-positive kinetochores per AP metaphase plate for day 27-30 organoids (e.g., D, E). Data are the mean ± SEM of ≥65 APs from five independent experiments, with a total of ≥7 organoids, for each of the two ESC lines. Bracket with ****, p <0.0001 (Mann-Whitney U test). (**G**) Mitotic APs (immuno)stained for FoxG1 (magenta) and DAPI (grey) in the indicated types of organoids. From left to right: from Figure 5D, this figure panel D, from Figure 5E, and this Figure panel E.

**Supplementary Figure 6.**
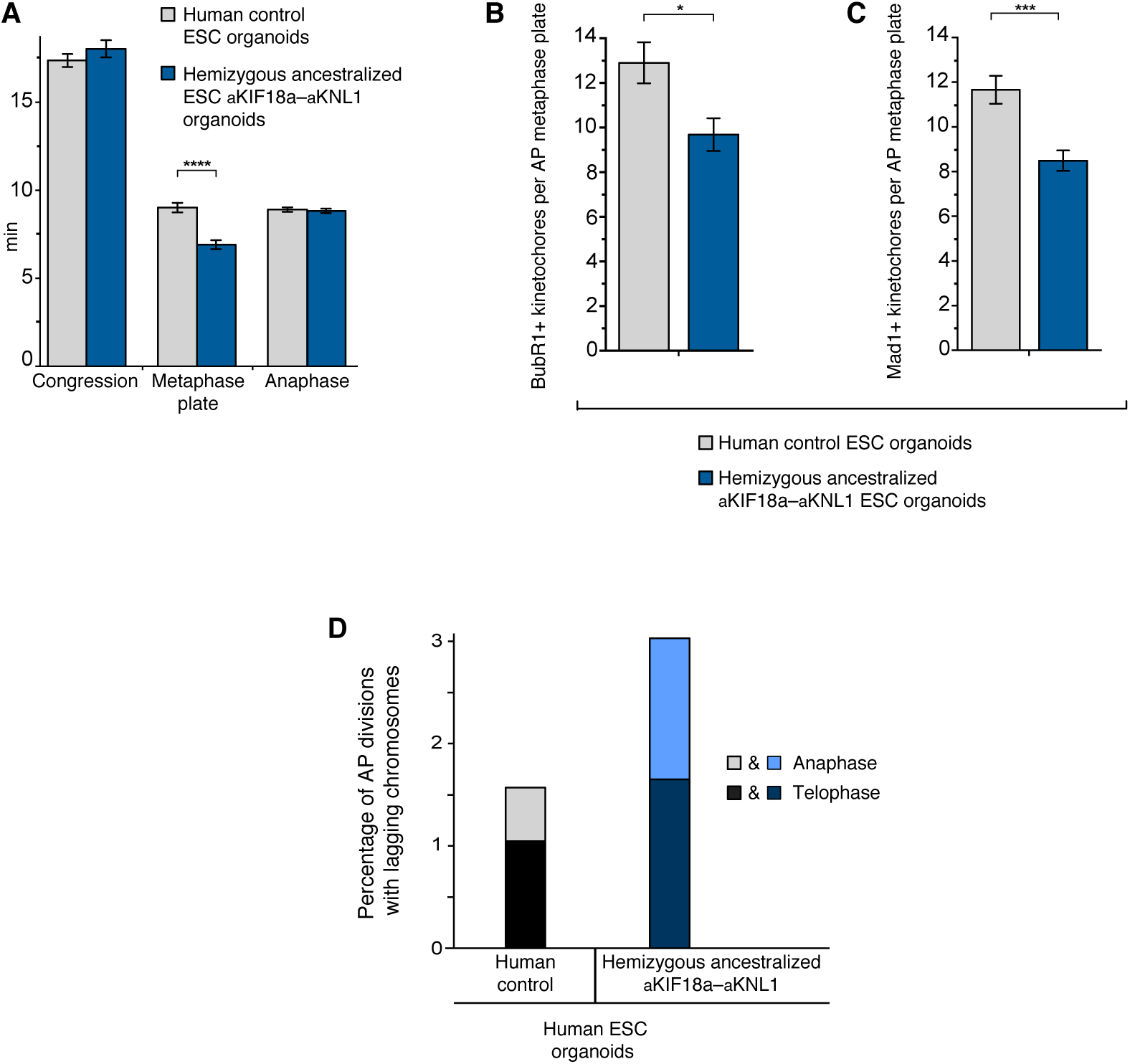
Organoids from ESC lines that are hemizygous for ancestralized KIF18a or KNL1. All panels show data similar to the panels indicated below, but with cerebral organoids from two H9 ESC lines, one where the gene *KIF18a* is hemizygous and edited to the ancestral, Neandertal-like state and the gene *KNL1* is homozygously edited, and the other where the gene *KNL1* is hemizygous and edited to the ancestral, Neandertal-like state and the gene *KIF18a* is homozygously edited. The results obtained with these two ESC lines were pooled. (**A**) Similar to Figure 4D. Times of “Congression”, “Metaphase plate” and “Anaphase” for APs in the two types of organoids. Data are the mean ± SEM of ≥134 APs from four independent experiments, with ≥7 organoids, for each of the lines. Bracket with ****, p <0.0001 (Mann-Whitney U test). (**B**) Similar to Figure 5F. Number of BubR1-positive kinetochores per AP metaphase plate in the two types of organoids. Data are the mean ± SEM of ≥35 APs from four independent experiments, with a total of ≥7 organoids, for each of the two lines. Bracket with *, p <0.05 (Mann-Whitney U test). (**C**) Similar to Supplementary Figure 4F. Number of Mad1-positive kinetochores per AP metaphase plate in the two types of organoids. Data are the mean ± SEM of ≥53 APs from five independent experiments, with a total of ≥7 organoids, for each of the two lines. Bracket with ***, p <0.001 (Mann-Whitney U test). (**D**) Similar to Figure 6I right. Percentages of AP divisions with lagging chromosomes in the two types of organoids. Data are the sum of ≥362 AP divisions from six experiments for each of the two types of organoids. The percentages for telophase (dark shade) and anaphase (light shade) are indicated separately.

### Supplementary Tables

**Supplementary Table 1.**
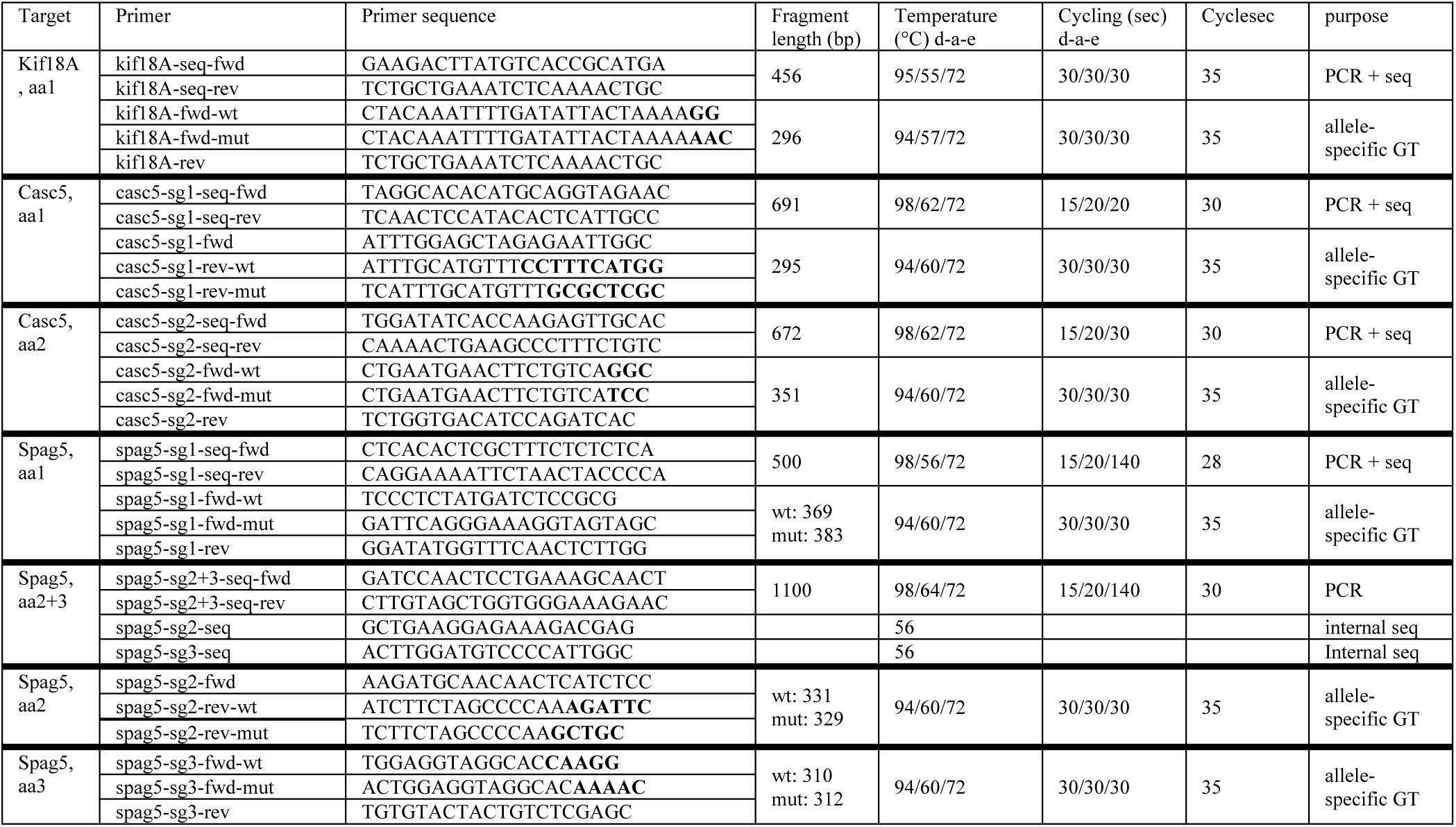
Overview of primers and sequencing conditions for genotyping of gene-edited mice. Primers used (a) for the sequencing of CRISPR/Cas9 events (PCR+seq); and (b) for allele-specific genotyping (GT) in wild type and humanized mice. Aa amino acid. Initial denaturation: 2 min at 95°C, d-a-e: denaturation – annealing – extension during PCR cycling, extension: 5 min at 72°C.

**Supplementary Table 2.** Quantification of *KIF18a*, *KNL* and *PPIB* mRNAs in cell line lysates. Numerical values correspond to cycle threshold in RT-qPCR; N/A indicates that there was no amplification of the targeted region.

| **Template** | ***Reverse transcriptase intact*** |  |  | ***Reverse transcriptase heat-inactivated*** |  |  |
| --- | --- | --- | --- | --- | --- | --- |
|  | ***KIF18a*** | ***KNL*** | ***PPIB*** | ***KIF18a*** | ***KNL*** | ***PPIB*** |
| RNA - ancestralized line 1 | 28.5 | 27.8 | 26.8 | N/A | N/A | N/A |
| RNA - ancestralized line 2 | 27.7 | 28.4 | 27.2 | N/A | N/A | N/A |
| RNA - non-edited line 1 | 27.1 | 27.9 | 26.7 | N/A | N/A | N/A |
| RNA - non-edited line 2 | 28.6 | 29.0 | 27.9 | N/A | N/A | N/A |
| H9-KR line | 27.5 | 28.1 | 27.1 | N/A | N/A | N/A |
| RNA extraction blank | N/A | N/A | N/A | N/A | N/A | N/A |
| DNA HeLa cells | N/A | N/A | N/A | N/A | N/A | N/A |
| water blank | N/A | N/A | N/A | N/A | N/A | N/A |

## Notes

### Competing Interest Statement

The authors have declared no competing interest.

